# Robust representation of natural images by sparse and variable population of active neurons in visual cortex

**DOI:** 10.1101/300863

**Authors:** Takashi Yoshida, Kenichi Ohki

## Abstract

Natural scenes sparsely activate neurons in the primary visual cortex (V1). However, how sparsely active neurons robustly represent natural images and how the information is optimally decoded from the representation have not been revealed. We reconstructed natural images from V1 activity in anaesthetized and awake mice. A single natural image was linearly decodable from a surprisingly small number of highly responsive neurons, and an additional use of remaining neurons even degraded the decoding. This representation was achieved by diverse receptive fields (RFs) of the small number of highly responsive neurons. Furthermore, these neurons reliably represented the image across trials, regardless of trial-to-trial response variability. The reliable representation was supported by multiple neurons with overlapping RFs. Based on our results, the diverse, partially overlapping RFs ensure sparse and reliable representation. We propose a new representation scheme in which information is reliably represented while the representing neuronal patterns change across trials and that collecting only the activity of highly responsive neurons is an optimal decoding strategy for the downstream neurons

## Introduction

Sensory information is thought to be represented by a relatively small number of active neurons in the sensory cortex. This sparse representation has been observed in several cortical areas^1-9^ and is postulated to reflect an efficient coding of the statistical features in sensory inputs^4, 10^. However, it has not been determined how small numbers of active neurons represent sensory information and how the information is optimally decoded from the sparse representation.

In the primary visual cortex (V1), a type of neuron termed the simple cell has a receptive field (RF) structure that is spatially localized, oriented, and has a bandpass filter property with a specific spatial frequency. This RF structure is modelled by a two-dimensional (2D) Gabor function^11^. According to theoretical studies, a single natural image is represented by a relatively small number of neurons using Gabor-like RFs, whereas information about multiple natural scenes is distributed across the neuronal population^10, 12, 13^. Indeed, V1 neurons respond sparsely to natural scenes at the single cell level^2, 3, 5-9^ and population level^3, 5, 14^. Population activity with higher sparseness exhibits greater discriminability between natural scenes^5^.

What types of information from natural scenes are represented in sparsely active neuronal populations in the brain? The visual contents of natural scenes or movies are reconstructed from single-unit activity in populations within the lateral geniculate nucleus (LGN) collected from several experiments^15^ and functional magnetic resonance imaging (fMRI) data from the visual cortices^16-19^. However, it has not been addressed experimentally whether the visual contents of natural images are represented by small numbers of sparsely active neurons and whether V1 RFs in the brain are useful in representing the natural image. It has also remained to be revealed which decoding strategy is optimal for the sparse representation. In the sparse representation, a single natural image activates a small number of neurons strongly and some remaining neurons weakly. Whether only a small number of strongly active neurons represent information, or the remaining neurons also have additional information is an important question. Furthermore, do the sparsely active neurons reliably represent the natural image contents, regardless of trial-to-trial response variability? Although a computational model^20^ has suggested that sparse and overcomplete representation is the optimal representation for natural images with unreliable neurons, this model has not been examined experimentally.

We also addressed how visual information is distributed among neurons in a local population. Some neurons are “unresponsive” to visual stimuli (e.g., the response rate of the mouse V1 to visual stimuli is 26–68%)^21-27^, indicating that only a subset of neurons represent sensory information. However, this may be partially attributed to incomplete coverage of stimulus properties on the RF properties of all neurons. Thus, there are two extreme possibilities: sparsely active neurons distributed among all neurons in a local population or only a specific subset of cells process the natural images. The proportion of neurons that are actually involved in information processing is a matter of debate^28, 29^.

Here, we examined how a small number of highly responsive V1 neurons represented natural image contents and how the information was optimally decoded from the sparse representation. Using two-photon Ca^2+^ imaging, we recorded visual responses to natural images from local populations of single neurons in the V1 of anaesthetized and passively viewing awake mice. A small number of neurons (a few percent) exhibited large responses to each natural image, which was sparser than that predicted by the linear encoding model. On the other hand, more than 80% of neurons were activated by at least one of the natural images, revealing that most neurons in a local population are involved in processing natural images. We reconstructed the natural images from the activity to estimate the information about the visual contents. The visual contents of a single natural image were linearly decodable from a small number of highly responsive neurons, and an additional use of remaining neurons even degraded the reconstruction performance. The highly responsive neurons showed diverse RFs, which helped small numbers of neurons represent complex natural images. Furthermore, the responsive neurons reliably represent the image, regardless of trial-to-trial response variability. The reliable image representation was supported by multiple neurons with partially overlapping representation. Further, neurons with overlapping representation were almost independently active across trials, which was beneficial to the reliable image representation across trials. Responsive neurons were only slightly overlapped between images, and many natural images were represented by the combinations of responsive neurons in a population. Finally, visual features encoded by neurons in a local population were sufficient to represent all the natural images used in the present study. These results revealed a new robust representation of a natural image by a small number of neurons in which information is reliably represented while the representing neuronal patterns change across trials and imply that collecting only the activity of highly responsive neurons is an optimal decoding strategy for the sparse representation. Preliminary results of this study have been published in an abstract form^30^ and on a preprint server^31^.

## Results

The main purpose of this study is to examine whether and how natural images are represented in the sparse representation scheme. We first confirmed the sparse response to natural images in our dataset. Next, we showed that the natural images were reconstructed from a relatively small number of responsive neurons. Finally, we addressed how the small number of neurons reliably represented natural images, regardless of trial-to-trial response variability.

### Sparse visual responses to natural images in anaesthetized mouse V1

We presented flashes of natural images as visual stimuli (200 images. Each image was consecutively flashed three times in a trial. Flash duration: 200 ms, flash interval: 200 ms. At least 12 trials for each image. Fig. 1a, see Methods). Using two-photon calcium (Ca^2+^) imaging, we simultaneously recorded the activity of several hundreds of single neurons from layers 2/3 and 4 of anaesthetized mouse V1 (560 [284–712] cells/plane, median [25–75^th^ percentiles], n = 24 planes from 14 mice, 260–450 microns in depth, see Fig. 1b for representative response traces).

**Figure 1.**
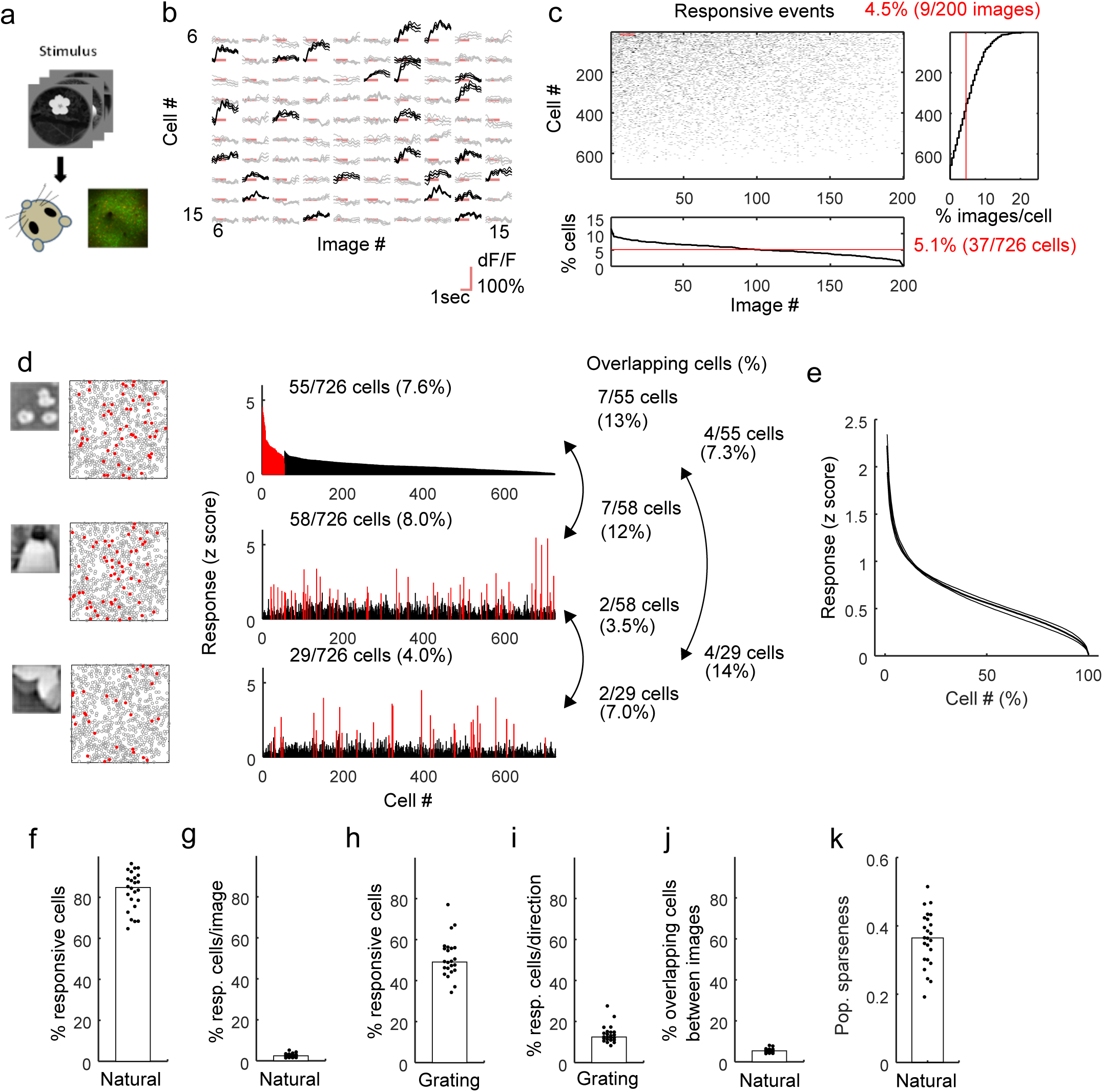
Sparse visual response to a natural image in mouse V1. **a**. Experimental schematic. Natural image flashes were presented as visual stimuli, and the activities of single neurons in the mouse V1 were recorded using two-photon Ca^2+^ imaging. **b**. Trial-averaged time courses of visual responses to 10 natural images (images #6–15 in a row) in 10 representative cells (cells# 6–15 in a column). The three lines for each response indicate the mean and the mean ± the standard error across trials. Black: significant responses, grey: non-significant response, red line: stimulus periods during which each image type was flashed three times. **c**. Plots of significant responses of all cells in an example plane (n = 726 cells, upper left panel). Responses shaded by the red line in the upper left panel correspond to responses presented in (b). The percentage of responsive cells for each image (bottom panel) and the percentage of images to which each cell responded (right panel) are shown as line graphs. Red lines (bottom and right panels) indicate median values. Cell numbers (cell #) were sorted by the percentage of images to which they responded, and image numbers (image #) were sorted by the percentages of cells that responded to each image in descending order. Single images activated relatively fewer neurons (bottom panel). **d**. Examples of population response patterns to three images. Left panels: Natural image stimuli and the spatial distributions of responsive cells in an imaging area (side length: 507 microns). The red-filled and grey open circles indicate the responsive and remaining cells, respectively. Right panels: Histograms of the visual responses of neurons in a local population. In the top panel, cells are divided into responsive (red bars) and the remaining groups (black bars) and are sorted by the response amplitude of each group to the natural image presented in the upper left panel (descending order). Visual responses to other images are plotted in the middle and bottom panels. The cell # order was fixed among the three histograms. Only a small number of responsive neurons are duplicated among the three images. **e**. Distribution of the amplitude of responses to single images. The cell # is sorted by the amplitude of the response to each image and averaged across images in a plane. After normalizing the cell # (x-axis), data were collected across planes (n = 24 planes). The median (thick line) and 25–75^th^ percentiles (thin lines) are shown. Small percentages of neurons exhibited higher response amplitudes. **f**. The percentages of visually responsive cells. Most cells in a population were visually responsive. **g**. The percentage of responsive cells per image. A small percentage of cells were responsive for each image. **h**. Percentages of responsive cells for the moving grating. **i**. Percentage of responsive cells for each direction of the moving grating. **j**. Percentages of overlap of responsive cells between the natural images. Only a small percentage of responsive cells overlapped between images, indicating that the cells responding to each image were distributed in a population. **k**. Population sparseness. **f–k**. Each dot indicates data obtained from one plane, and the medians of 24 planes are shown as bars

We first found that most neurons in a local population were visually responsive. Responsive neurons were determined by one-way analysis of variance (ANOVA, p < 0.01, data with responses to 200 images and one baseline in each trial, see Methods). Across planes, the percentage of responsive neurons was 85% [77–90%] (Fig. 1f). We also estimated a false positive rate of this criterion with label-shuffled data and found a 1.0% [0.9–1.0%] false positive rate. Thus, most neurons were involved in visual processing.

We next examined what percentage of neurons responded to each image. For each responsive neuron determined by the ANOVA, a significant response to each image was defined by a t-test and amplitude threshold (p < 0.01 using a t-test and >10% trial-averaged response change, activity during stimulus vs. baseline just before the stimulus) (see Methods and Supplementary Fig. 1). Fig. 1c presents plots of significant visual response events for all images (x-axis) across all neurons (y-axis) in an example plane (n = 726 cells, depth: 360 microns from the brain surface). Hereafter, we call these significant responses highly responsive. In the example plane, each responsive neuron responded to 4.5% (9/200) images (Fig. 1c, right panel), and a few to 10% of neurons were highly responsive to a single image (5.1% (37/726) cells/image, Fig. 1c, bottom panel), indicating sparse visual responses to natural images. Across planes, the percentage of responsive neurons for each image was 2.5% [1.8–3.0%] (Fig. 1g, Supplementary fig. 1b). Thus, only a few percent of neurons were responsive for each image. This low response rate to each image was not due to poor recording conditions. The same neurons responded well to moving gratings (The percentage of responsive cells for at least one direction: 49% [45–56%]. The percentages of responsive cells for each direction of grating: 12.5% [11–14.8%], Fig. 1h and 1i).

The highly responsive neurons only slightly overlapped between images. Fig. 1d presents representative activity patterns for three natural images (Fig. 1d, left column). Each image activated different subsets of neurons that exhibited small overlaps between images (Fig. 1d, right column). Of the responsive cells, 5.4% exhibited overlap between two images (25–75^th^ percentiles for 24 planes: 4.8–6.0%, Fig. 1j). We further computed the distributions of the response amplitudes to single images (Fig. 1e). Only a small number of neurons exhibited visual responses with greater amplitudes, which is a property of a sparse representation (Fig. 1e). The population sparseness^2, 3^, a measure of a sparse representation, was comparable to that of a previous report for mouse V1^5^ (0.36 [0.30–0.42], Fig. 1k, see Methods). Thus, each natural image activated a relatively small number of neurons, whereas most neurons in a local population were visually responsive, suggesting the sparsely distributed representation of natural images in V1 that was originally proposed in a previous study^10^. The latter result also represents the first report of the visual responsiveness of most neurons in mouse V1 to natural image stimuli^28, 29^.

### Partially overlapping representations of visual features among local V1 populations

We created an encoding model for the visual responses of an individual neuron to examine visual features represented by each neuron. We used a set of Gabor wavelet filters (1248 filters, Fig. 2a and 2b) to extract the visual features from the natural images. Down sampled natural images (**I**) were applied to Gabor filter (**G**_fwd_) and were transformed into sets of feature values (Gabor feature values, **F**).

**Figure 2.**
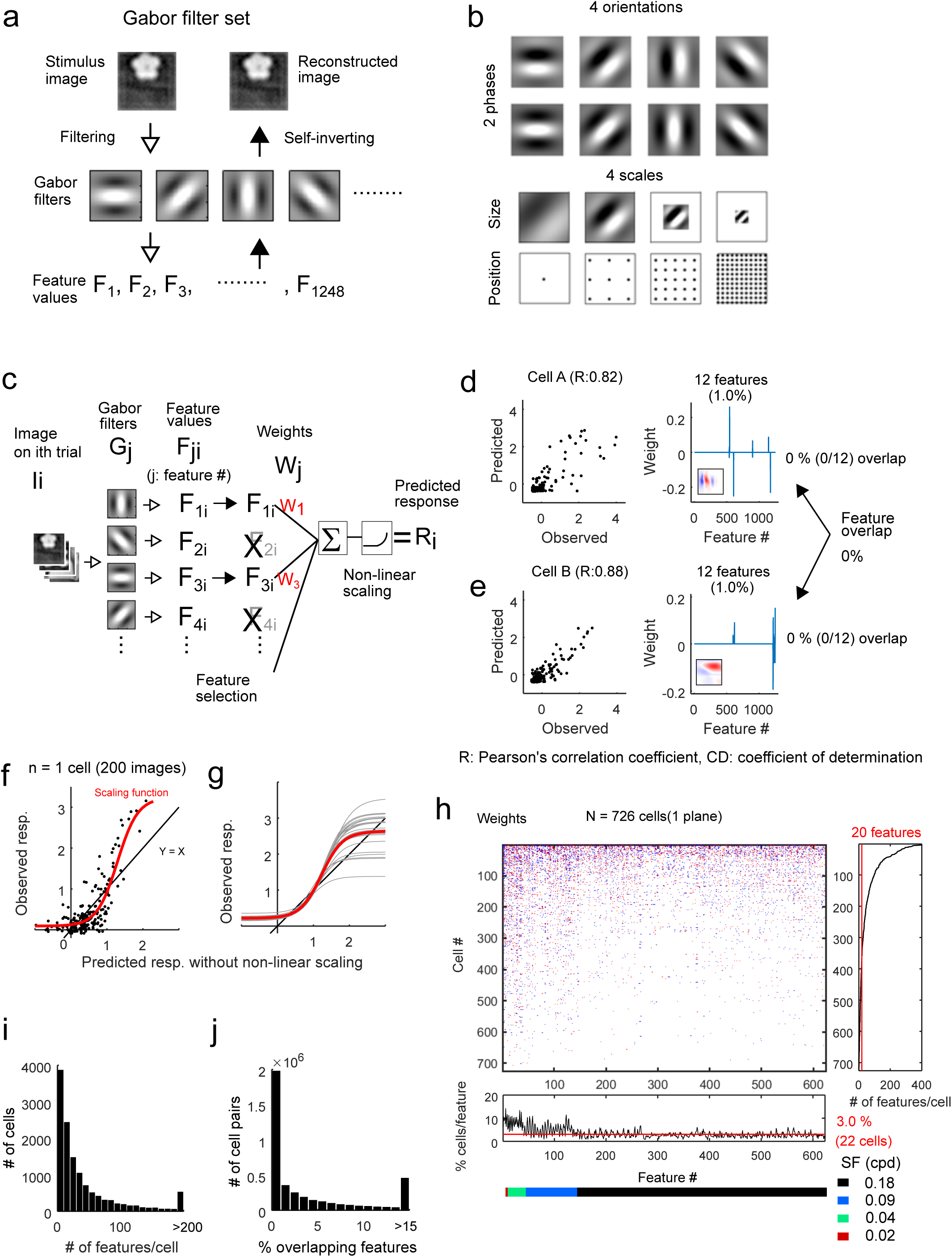
Small overlap in the encoded visual features among cells in a local population. **a**. Schematic of transformation between natural image and feature values with Gabor filters. Each natural image was subjected to Gabor filters to obtain the corresponding Gabor feature values. Conversely, a set of Gabor feature values were transformed into an image by summing the Gabor filters after multiplying by the corresponding Gabor feature values. **b**. Schematics of the Gabor filter set. Four orientations, two phases, four scales (or spatial frequencies) of Gabor filters were used. The four scales of Gabor filters (spatial frequency: 0.02, 0.04, 0.09, and 0.18 cycle per degree) were positioned on 1 × 1, 3 × 3, 5 × 5, and 11 × 11 grids. A total of 1248 filters were used. **c**. Schematic of the encoding model of the visual response of a single cell. The visual response is represented by weighted sum of the selected Gabor feature values obtained from a set of Gabor filters in each cell. The predicted visual response to the image in the ith trial (R_i_) is represented by the following equation: R_i_ = NL(∑W_j_×F_ji_+b), where NL(·) is a non-linear scaling function, W_j_ is the weight for jth Gabor feature, and F_ji_ is the feature value for the jth Gabor filter (Gj) obtained from ith trial image (Ii). The Gabor feature was selected based on the correlation between its feature values and visual response (Feature selection, see Methods). **d**. and **e**. Representative predicted responses of two neurons. Left panels: Comparison between the observed and predicted responses, respectively. Each dot indicates a response to one image. Right panels: Weight parameters of the representative neurons presented in the left panels. The weights of one of 10 models (each model corresponds to one of the 10-fold cross validations) are shown. The number of non-zero weights (i.e., number of used features) is shown above the panels. Encoding filters (weighted sums of Gabor filters) are shown in the insets (red and blue indicate positive and negative values, respectively). **f**. Comparison of the response predicted by only the linear step (regression of Gabor feature values without non-linear (NL) scaling) and the observed response in the representative neuron shown in Fig. 2**e**. Each dot indicates a response to one image. The red curve indicates the NL scaling function curve (see Methods). The NL step resulted in an enhancement of the sparse visual responses. The black line indicates y = x line. **g**. NL scaling function curves across planes. Each grey curve was obtained by averaging the NL scaling curves across cells in each plane. Red curve indicates the averaged curve across planes (n = 24 planes). The black line indicates y = x line. **h**. Upper left panel: Raster plot of the weights in the plane illustrated in Fig. 1c (red: positive weight, blue: negative weight). The median values for the models of the 10-fold CVs are shown. Right panel: Percentage of features used for each cell. Bottom panel: Percentage of cells in which each feature was involved in the response prediction. The coloured bar under the x-axis indicates the spatial frequency of the Gabor filter corresponding to each feature. Red lines in the bottom and right panels indicate median values. Only half of the Gabor features (624/1248 with one of two phase sets) are shown for visibility, but the remaining features were included in the data shown in the right panel. SF: spatial frequency. **i**. Distribution of the number of features used in each cell (n = 12,755 cells across 24 planes). **j**. Distribution of the percentages of features that overlapped between cells (n = 3,993,653 cell pairs across 24 planes).

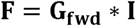

For each neuron, we first selected the Gabor features that exhibited strong correlations with the visual response. The correlation threshold for the selected feature was adjusted to maximize the visual response prediction in each neuron (Fig. 2c. Supplementary Fig. 2a and 2b). Then, the visual responses of a single cell (**R**^k^, the kth cell’s responses) were represented by a linear regression of the selected feature values (**F**_select_) followed by non-linear scaling (NL( ), Fig. 2c, see Methods).

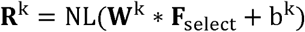

Parameters of the regression model (weights, **W**^k^ and a bias, b^k^) were estimated with 90% of the dataset, and the response prediction performance of the model was estimated with the remaining 10% dataset (10-fold cross-validation (CV. **W**^k^ and b^k^ were estimated in each CV). The model was obtained independently for each cell (i.e., for each k).

The results of two example neurons are shown in Fig. 2d and 2e. The response prediction performances (the correlation coefficients between the observed responses and the responses predicted by the model) were 0.82 and 0.88. These neurons were represented by 12 (out of 1248) Gabor features (Fig. 2d and 2d, right panels), and their encoding filters (weighted sums of the Gabor filters) were spatially localized (Fig. 2d and 2d, insets in the right panels).

The median prediction performance of the encoding model was 0.32 in the example plane presented in Fig. 1 (25–75^th^ percentiles: 0.15–0.50, n = 726 cells) and 0.21 in all cells across planes (25–75^th^ percentiles: 0.06–0.42, n = 12,755 cells across 24 planes, Supplementary Fig. 2d). The non-linear scaling step suppressed weak predicted responses and enhanced strong predicted responses (Fig. 2f and 2g), suggesting that this non-linear step enhanced the sparseness of the predicted response obtained from the linear step (i.e., linear regression of feature values). The non-linear step also slightly enhanced the performance (Supplementary Fig. 2c).

The visual response of an individual neuron was represented by a small number of Gabor features. On average, each neuron encoded 20 (out of 1248) features in the example plane (25–75^th^ percentiles: 8–52. Fig. 2h) and 21 features in all recorded neurons (25–75^th^ percentiles: 8–51, n = 12,755 cells, Fig. 2ig and Supplementary Fig. 2e). The features encoded by each neuron were spatially localized and had similar orientations (Supplementary Fig. 3a–d). These features were also related to the RF structure of each neuron. The RF structure of each neuron was estimated using the regularized inverse method^32-34^ (see Methods). The regression weights of the Gabor features in the encoding model were positively correlated with the similarity between the corresponding Gabor filter and the RF structure (Supplementary Fig. 3e–3h).

**Figure 3.**
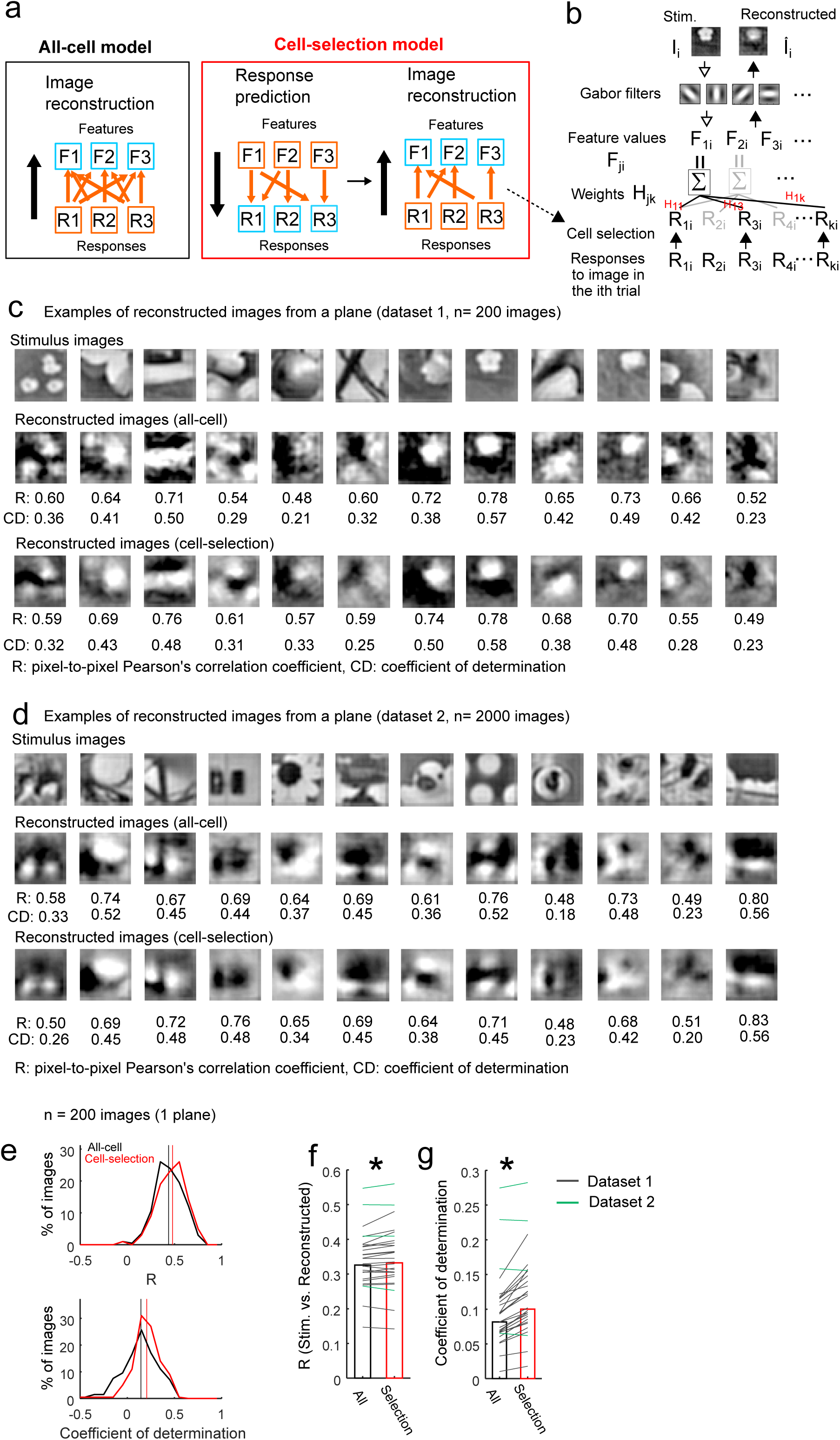
Image reconstruction based on population activity. **a** and **b**. Schematic of the image reconstruction models. **a**. In the image reconstruction, each feature value was reconstructed using responses of multiple cells. In the all-cell model, each feature value was reconstructed by all cells. In the cell-selection model, the feature value was reconstructed by selected cells for each feature. The cell selection was based on the response prediction model for each cell; each cell participates in the reconstruction of features that the cell encodes in the response prediction model. **b**. Details of the image reconstruction model. For each Gabor feature, j, feature values (F_ji,_ i: trial # across stimuli and trials, j: Gabor feature #) were independently regressed (weights: H_jk_, k: cell #) by multiple cell responses (R_ki_) to the image (Ii) in the ith trial (F_ji_ = ∑(H_jk_ × R_ki_) + c. c: a bias, not shown in the figure). Then, a set of reconstructed features (F_1i_, F_2i_, …., F_ji_) was transformed into an image (Î_i_). The flow of the reconstruction model is represented by black arrows from the bottom to the top. **c**. Examples of reconstructed images from main datasets (dataset 1, 200 images). Stimulus images (top), images that were reconstructed using the all-cell model (all-cell, middle) and using the cell-selection model (cell-selection, bottom) are shown. Each reconstructed image was averaged across trials. The reconstruction performances (R and CD) were computed for each trial, and trial-averaged performances are presented below each reconstructed image. **d**. Examples of reconstructed images from other datasets (dataset 2, 1000–2000 images). **e**. Distributions of R (top) and CD values (bottom) for the all-cell model (black lines) and the cell-selection model (red lines) in the example plane shown in Figs. 1 and 2 (n = 200 images reconstructed using 726 cells from a plane). Vertical lines indicate median values. **f** and **g**. R (f) and CD (g) of dataset 1 (black lines and bars) and of dataset 2 (green lines) across planes. *: p = 0.006 in (f) and p = 1.8×10^−5^ in (g) using the signed-rank test (n = 24 planes for dataset 1, and n = 4 planes for dataset 2). The reconstruction performance of the cell-selection model was comparable with or slightly higher than that of the all-cell model.

The Gabor features encoded by one neuron partially overlapped with those of other neurons (Fig. 2j). In the example neuros (Fig. 2d and 2e), none of the features overlapped. For all neuron pairs, the median overlap was 1.0% (25–75^th^ percentile: 0.0–5.6% relative to features represented by each cell, Fig. 2j and Supplementary Fig. 2f). The feature overlap between neurons was positively correlated with the similarity of the forward filter structure (Supplementary Fig. 3i and j). Furthermore, the diverse features encoded by individual neurons were also reflected in the distribution of forward filter similarity between cells (correlation coefficient between reverse filters: 0.0 [-0.03–0.04], Supplementary Fig. 3k). Based on these findings, the Gabor features encoded by individual neurons in a local population were highly diverse and partially overlapped.

The analysis of the encoding model also revealed how the individual Gabor features were encoded across neurons (upper left and bottom panels in Fig. 2h). As the spatial frequency (SF) of the Gabor filter increased (i.e. the scale decreased), the corresponding feature contributed to the visual responses of fewer neurons (Fig. 2h bottom). This pattern likely reflected the fact that Gabor filters with a low SF (i.e., a large scale) covered more of the neuron’s RF, whereas Gabor filters with a high SF (i.e., a small scale) affected the responses of fewer neurons. Furthermore, almost all features contributed to the responses of at least one cell (Supplementary Fig. 2g).

### Image reconstruction from the activity of the neuronal population

We next examined whether the features encoded in a local population of neurons represent the visual contents of the natural images. To examine this, we reconstructed stimulus images from the neuronal activity^15-19^. In the image reconstruction model, each Gabor feature value (**F**^j^, the jth Gabor feature values) was subjected to a linear regression analysis of the activity data of multiple neurons (**R**) with model parameters of weights (**H**^**j**^) and a bias (c^j^) (Fig. 3a and b).

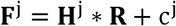

Each Gabor feature value was independently reconstructed (i.e., **H**^**j**^ and c^j^ were estimated for each feature, j). Then, the sets of reconstructed feature values 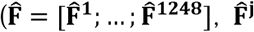: reconstructed jth feature values) were transformed into images (**I**) using Gabor filter for image reconstruction (**G**_rev_ Fig. 3b, see Methods).

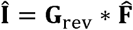

The reconstruction performance was estimated with a different test dataset from the training dataset used in the regression analysis (10-fold CV with the same data split to that in the encoding model. **H**^**j**^ and c^j^ were estimated in each CV).

We first used a model in which each feature value was reconstructed from all neurons (all-cell model, Fig. 3a). In the example plane (n = 726 neurons, presented in Figs. 1 and 2), the rough structures of the stimulus images were reconstructed from the population activity (Fig. 3c). The pixel-to-pixel correlation between stimulus and reconstructed images (similarity of image patterns) was 0.43 [0.35–0.55] (median [25–75^th^ percentiles] of 200 images, dataset 1) in the example plane (n = 726 cells, Fig. 3e upper panel) and 0.33 [0.30–0.37] across all planes (n = 24 planes, Fig. 3f). The coefficient of determination (CD, goodness of model prediction, see Methods) was 0.14 [0.02–0.26] in the example plane and 0.08 [0.07–0.1] across planes (Fig. 3e bottom and 3g). We further confirmed that our image reconstruction could be used for another dataset (dataset 2, 1000–2000 images that did not include original 200 images). The performances of dataset 2 were similar to those of the original dataset (R: 0.45 [0.34–0.52], CD: 0.19 [0.11–0.25], n = 4 planes from three mice. Fig. 3d, green lines in Fig. 3f and 3g). Thus, the visual contents of natural images were extracted linearly from the neuronal activity of the local population.

We next used another reconstruction model in which each feature was reconstructed from a subset of neurons that showed strong correlations between the feature values and responses (cell-selection model, Fig. 3a). In the cell-selection model, each neuron participated in reconstructions of subsets of features that the neuron encoded in the response prediction model (Fig. 3b, see Methods). The reconstruction performances of the cell-selection model were almost comparable or even slightly higher than those of the all-cell model (Dataset 1, R: 0.33 [0.3–0.38]. P = 0.006 by signed rank test compared to all-cell model. CD: 0.10 [0.08–0.14]. P = 1.8×10^−5^ by signed rank test. N = 24 planes. Dataset 2, R: 0.45 [0.33–0.53], P = 0.9 by signed rank test compared to all-cell model. CD: 0.19 [0.11–0.25], P = 0.9 by signed rank test. N = 4 planes. Fig.3c–g). The performances were not substantially affected by the cell-selection step using nested CV (Supplementary fig. 4, see Methods).

**Figure 4.**
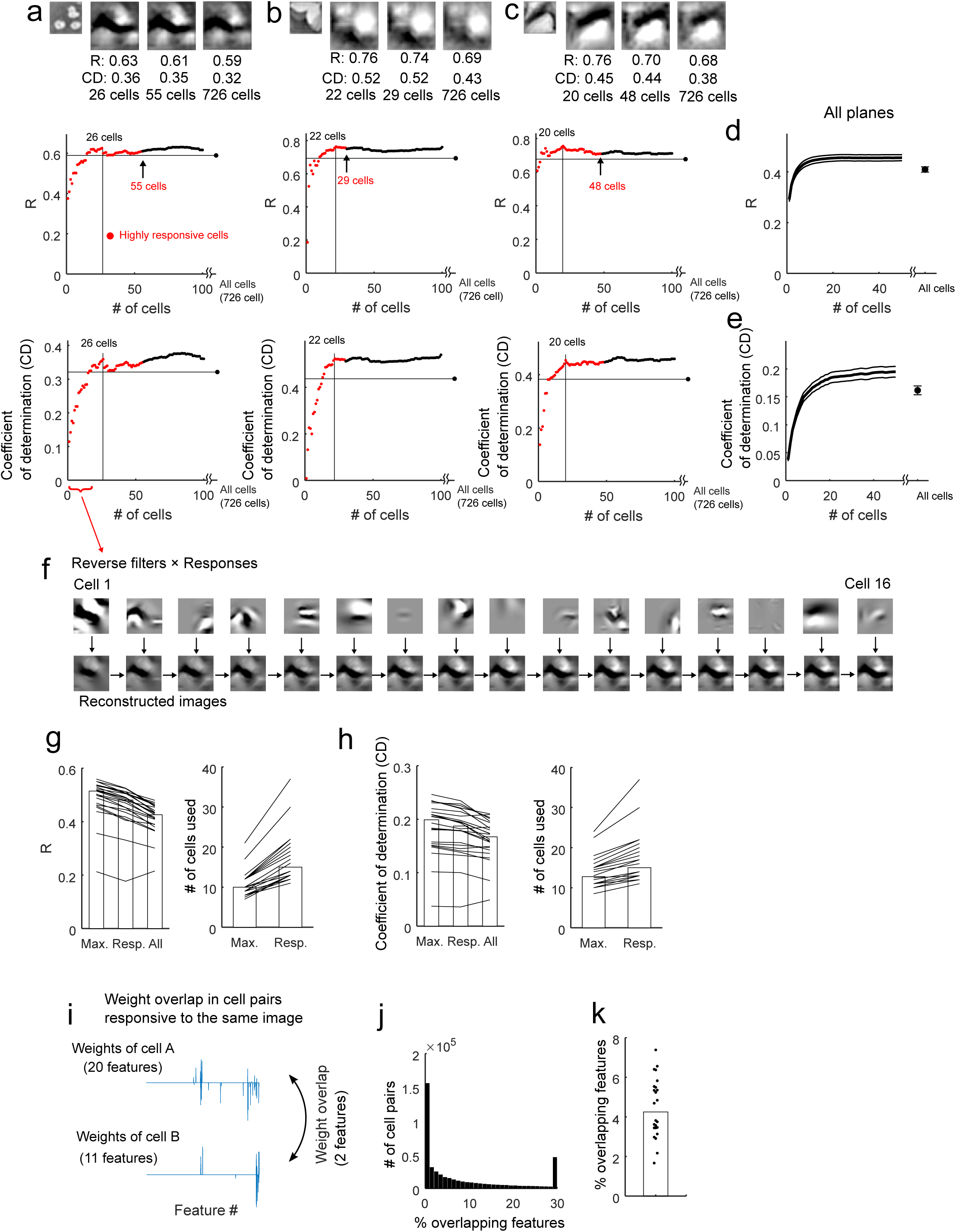
A small number of highly responsive neurons mainly represent the visual contents of a single natural image. **a–c**. Top panels: Examples of reconstructed images from a subset of highly responsive cells and all cells. Stimulus images (1^st^ panel) and reconstructed images (trial-average) from a subset of or all cells (2^nd^–4^th^ panels) are shown. Middle and bottom panels: Reconstruction performances (middle, R; bottom, CD) plotted against the number of cells used for the reconstructions. Using the weight parameters of the cell-selection model, the number of cells used to reconstruct each image was successively increased. The cells were first collected from the responsive cells (red dots) and then from the remaining cells (black dots), and the cells were ordered in each group using evoked response amplitude (descending order). The horizontal lines indicate the performance from all cells, and the numbers of cells for peak performance within the number of responsive cells are indicated by the vertical lines. In each case, the reconstruction performances of responsive cells were similar or slightly higher than those of all cells. **d** and **e**. Average performance curve (d, R; e, CD) plotted against the cell number. The thick black line and grey lines indicate the means and the means ± standard errors, respectively (n = 24 planes). Data for images including at least 10 responsive cells were only used. **f**. The contributions of the top 16 responsive cells to the image reconstruction shown in (a). In the top panels, reverse filters (weighted sum of Gabor filters) multiplied by the visual responses reveal the spatial patterns of an individual cell’s contributions to the reconstructed image. The patterns vary among cells. In the bottom panels, the reconstructed image gradually changes with the consecutive addition of single cells. **g** and **h**. Left panels: Performances (g, R; h, CD) obtained from all cells (All), highly responsive cells (Resp.) and peak performance (Max.) were compared. The peak performance was detected within the number of responsive neurons. Right panels: The number of cells used for the reconstruction was compared. Peak performances tended to be obtained from fewer cells than the number of responsive cells. (g, left) P = 5.4×10^−5^ for Max. vs. Resp.; P = 1.2×10^−4^ for Resp. vs. All; P = 6.2×10^−5^ for Max. vs. All. (g, right) P = 5.4×10^−5^. (h, left) P = 5.4×10^−5^ for Max. vs. Resp.; P = 3.3×10^−4^ for Resp. vs. All; P = 1.3×10^−4^ for Max. vs. All. (h, right) P = 1.7×10^−5^ using the signed-rank test (n = 24 planes). Each line indicates data for each plane, and bars indicate medians. In the analyses (**g** and **h**), we used only data for images that had at least 10 responsive cells. **i–k**. Overlap of weights (i.e., features) between the cells that were highly responsive to the same image. (**i**) Schematic of the analysis. (**j**) Distribution of the percentages of overlapping features in all cell pairs responding to the same image collected across planes. (**k**) The median percentages of overlapping features in the cell pairs responding to the same image. Each dot indicates the median in each plane (n = 24 planes) and the bar indicates the median across planes. The percentages of overlapping features were still low even in the cell pairs responding to the same image.

In the cell-selection model, each neuron participated in the reconstructions of a small number of features that each neuron encoded (Fig. 3b; also see Fig. 2 for encoding model analyses). We examined how the reconstruction performance was affected by an increase or decrease in the number of features for which each neuron participated in the reconstruction and found that the original model demonstrated nearly optimal performances (Supplementary Fig. 5a–c). Thus, how the individual neurons are involved in the reconstruction of Gabor features is one of the important factors of the image reconstruction, and the cell-selection model captures nearly optimal feature-cell assignment for the reconstruction.

**Figure 5.**
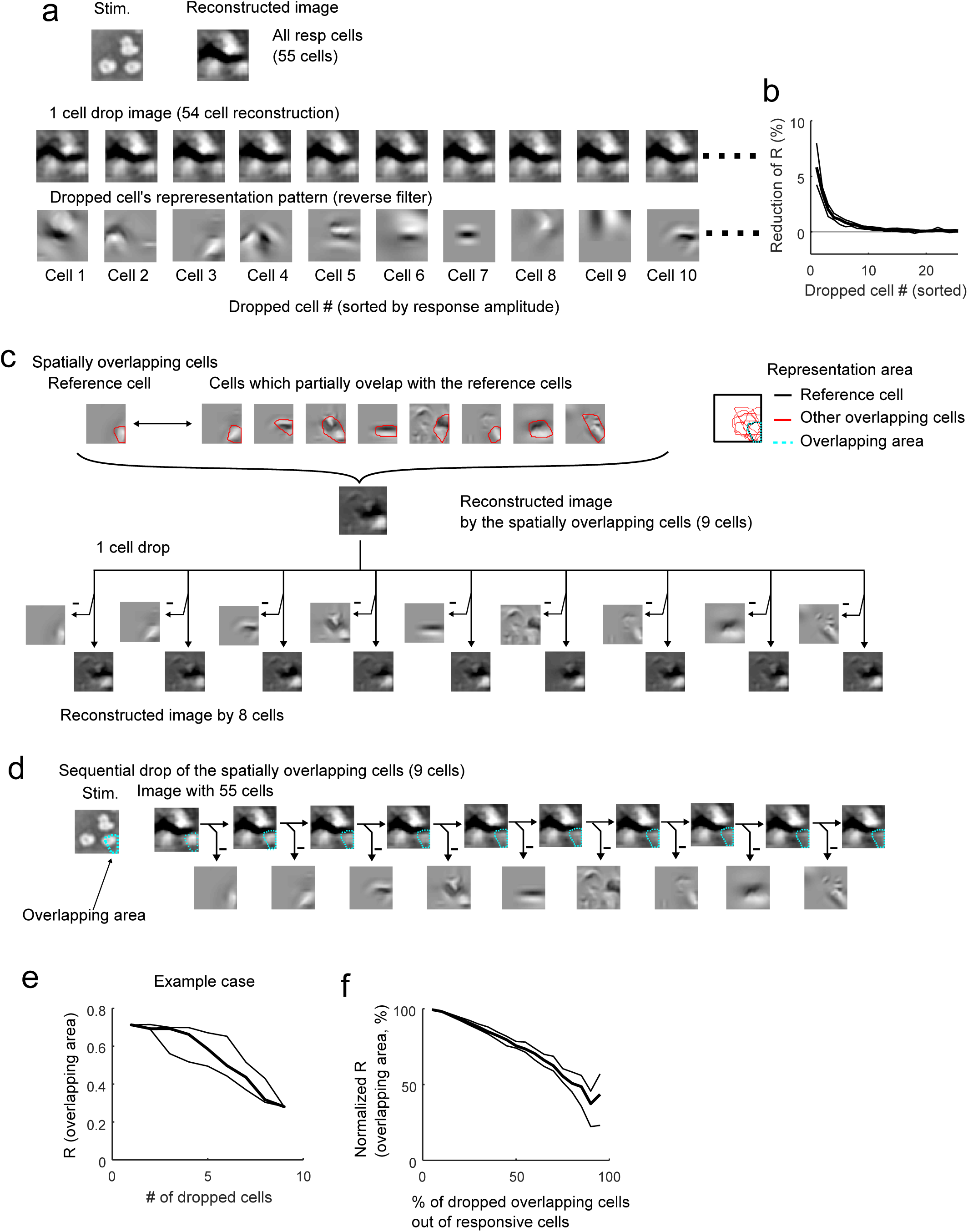
Robust image representation by small numbers of responsive neurons. **a**. Representative reconstructed images after the drop of a single cell. Top panels: Stimulus and reconstructed image obtained from all responsive neurons (55 cells). Middle panels: Reconstructed images obtained after the drop of a single cell. Bottom panels: Representation patterns (reverse filters) of the dropped cells. The cell number (cell #) is the same as shown in Fig. 4f. **b**. Reduction in reconstruction performance after removing a single cell. The cell # on the x-axis were ordered from largest to smallest response amplitudes. The cell # was the same order as shown in Fig. 4d and 4e. Thick line: median. Thin lines: 25^th^ and 75^th^ percentiles. N = 24 planes. **c**. Top panels: Reverse filters of overlapping cells (Example nine cells). The representation area of each neuron was contoured by the red line and overlaid on the right panel. Middle panel: Reconstructed image obtained from the nine overlapping cells. Bottom panels: Reconstructed images obtained from single cells (upper panels) and reconstructed images after the drop of a single cell (lower panels). The drop of a single cell exerted only a small effect on the reconstructed images. **d**. Representative reconstructed images obtained during the sequential drop of the nine overlapping neurons. Cyan dotted lines indicate the overlapping area of the nine cells. The quality of the reconstructed image around the overlapping areas gradually degraded after the drop of each cell. **e** and **f**. Plot of the R (or normalized R) for a local part of the reconstructed image (overlapping area) against the number (or percentage) of dropped cells for the representative case shown in Fig. 5c (e) and for summary of all data (f). Data were collected and averaged across cells and across stimuli in each plane and then collected across planes. Thick lines: medians. Thin lines: 25^th^ and 75^th^ percentiles obtained across repetitions of random drop (n = 120 repetitions, e) or across planes (n = 24 planes, f).

We also examined the image reconstruction performance in each spatial frequency component (0.02–0.18 cycle/degrees, cpd). Performances of lower spatial frequencies (0.02 and 0.04 cpd) were better than those of higher spatial frequencies (0.09 and 0.18 cpd) (R: 0.65, 0.53, 0.18, and 0.04, and CD: 0.24, 0.23, 0.02, and -0.03 for 0.02, 0.04, 0.09, and 0.18 cpd, respectively. Supplementary Fig. 5d–f). This may reflect the spatial frequency preference of mouse V1 neurons^35^. V1 neurons in a local population mainly represented information about the low-spatial-frequency components of the natural image.

### The visual contents of natural images are linearly decodable from small numbers of responsive neurons

A single natural image strongly activated a small number of neurons (Fig. 1). We next examined whether the small number of highly responsive neurons mainly represents a single image or the remaining neurons also contained information about the image. For this purpose, we changed the number of neurons used in the reconstruction of each image and examined how the reconstruction performance was affected. In this image reconstruction from a subset of cells, the weights of image reconstruction model were estimated based on the weights of the cell-selection model (See equation 5 in Methods). The weights were separately computed for each subset of cells, and a different set of cells was used for each image.

The results of the example plane are presented in Fig. 4a–4c. In each image, neurons were sorted by visual response amplitude (descending order) first among the responsive neurons (red dots in Fig. 4a–4c) and then among the remaining neurons (black dots in Fig. 4a–4c). The image was reconstructed by the top N neurons (N = 1–726 cells), and the reconstruction performances were plotted against the number of neurons used (Fig. 4a–4d). Approximately 20 neurons were sufficient to reconstruct the images with a level of peak performance, on average (Fig. 4d and 4e). The reconstruction performances of highly responsive neurons were higher than those of all cells (left panels in Fig. 4g and 4h). Furthermore, when we searched the peak performance within the number of responsive neurons in each image, the peak performances were obtained with fewer neurons than the highly responsive neurons (right panels in Fig. 4g and h). Based on these results, highly responsive neurons mainly represent the image, and the additional use of remaining neurons even decreases the reconstruction performance.

The features represented by individual neurons should be diverse to represent a natural image using a small number of neurons. Fig. 4f illustrates how individual responsive neurons contributed to image reconstruction in the case presented in Fig. 4a. Each neuron had a specific pattern of contributions (reverse filter: sum of Gabor filters × weights, see Methods), and the patterns varied across neurons (Fig. 4f, top panels) while partially overlapping in the visual field. In neuron pairs that were responsive to the same image, the number of overlapping Gabor features slightly increased compared to all pairs, but the percentage of overlapping features was still less than 5% (Responsive cell pair: 3.2% [0–13%] of features for all pairs and 4.3% [3.4–5.5%] across planes. All cell pairs: 1.0% [0–5.6%] for all pairs and 1.0% [0.9–1.2%] across planes, Fig. 4i–4k, cf. Fig. 2j, Supplementary Fig. 2f). These small overlaps and the diversity in the represented features among neurons should be useful for the representation of a natural image by a relatively small number of responsive neurons.

### Robust image representation by neurons with spatially overlapping representation

We next examined whether a single image was robustly represented by a small number of responsive neurons. We computed the reconstruction performance after dropping individual responsive cells (Fig. 5a and 5b; The cell # on the x-axis is the same as in Fig. 4d). The drop of a single cell exerted only a small effect on the reconstructed image (middle panels in Fig. 5a). A reduction of approximately 5% in reconstruction performance was observed for the best-responding neurons, and most neurons did not exhibit a change in performance (Fig. 5b). Thus, an image was robustly represented by highly responsive neurons against a single-cell drop.

This robustness against cell drop was due to the spatial overlap of representation (i.e., reverse filters) among responsive neurons (Fig. 5c). To analyse this point, we selected a set of overlapping cells for each responsive neuron; the overlapping cells consisted of one responsive neuron as a reference (cell 1) and other responsive neurons whose representation areas partially overlapped with that of the reference cell (top panels in Fig. 5c and Supplementary Fig. 6a-d, see Methods). Then, we examined the effects of cell drop on the reconstruction of a local part of the image from the overlapping cells. Although a single-cell drop had almost no effect on the reconstructed local image (bottom panels in Fig. 5c), a sequential drop of these cells gradually degraded the part of the reconstructed image (Fig. 5d). The pixel-to-pixel correlation between the stimulus and the reconstructed image in the overlapping area gradually decreased as the number of dropped cells increased (Fig. 5e and 5f). Based on these results, the robust image representation was due to neurons with spatially overlapping representations.

**Figure 6.**
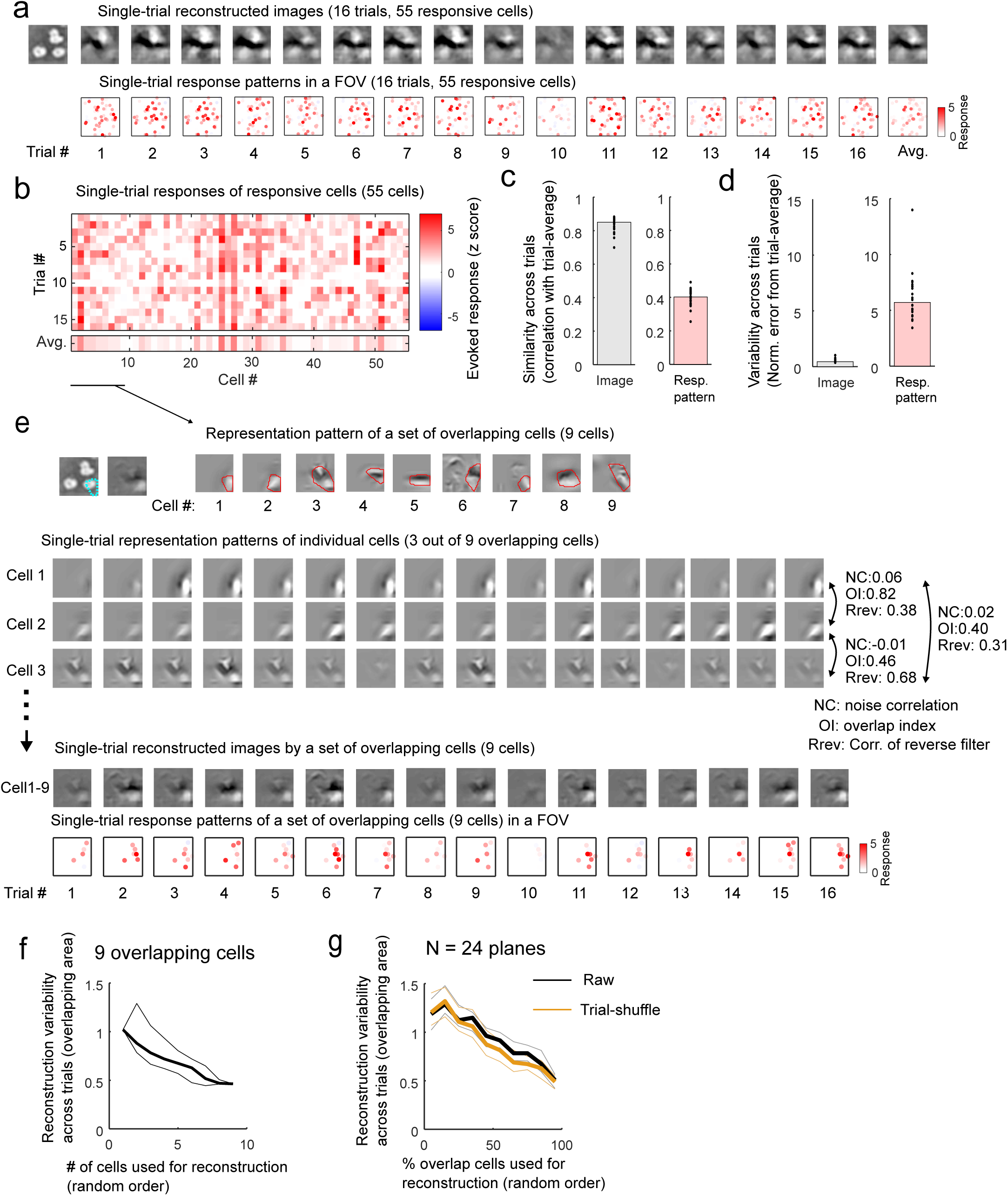
Reliable image representation across trials. **a**. Examples of single-trial reconstructed images (top panels) and response patterns in a FOV (bottom panels). The first panel is a stimulus image, and the last panel is a trial-averaged image. FOV size: 507 micron each side. Colour code for each dot indicates response amplitude of each cell. **b**. Single-trial evoked responses to the image in (**a**). **c**. Across-trial similarity of reconstructed images (left) and response patterns of responsive cells (right). N = 24 planes. The across-trial similarity was a Pearson’s correlation between a single-trial reconstructed image (or responses) and trial-averaged image (or responses). **d**. Across-trial variability of reconstructed images (left) and response patterns of responsive cells (right). N = 24 planes. Normalised squared error between single-trial image (or response patterns) and trial-averaged image (or response patterns) was computed and used for the across-trial variability. **e**. Reconstructed image from a set of overlapping cells (Cell# 1–9 in **b**). Upper left panels: Stimulus image and trial-averaged reconstructed images from the nine overlapping cells. The cyan dotted line indicates the overlapping area. Upper right panels: Representation (reverse filters) of the overlapping cells. The red line in each panel indicates the representation area. Cell# 1 was the reference cell. Lower panels: Single-trial representation patterns of three example cells (cell 1, 2, and 3) selected from the overlapping cells. Bottom panels: Single-trial reconstructed images (upper) and single-trial response patterns in a FOV (lower) obtained from the nine overlapping cells. Brightness of each colour dot in lower panels indicates response amplitude of each cell. **f**. Across-trial variability of a local part of reconstructed image (cyan dotted line in **e**) against the number of overlapping cells used for the reconstruction in the example case in **e**. Thick and thin lines were median and 25^th^ or 75^th^ percentile of the variability among 200 random sequences of cell adding. **g**. Across-trial variability against the percentage of the overlapping cells used for the reconstruction. N = 24 planes. Black lines: Raw data. Orange lines: trial-shuffled data. Thick and thin lines were median and 25^th^ or 75^th^ percentile of the variability among 24 planes.

### Overlapping representation provides a reliable image representation, regardless of trial-to-trial variability

We further analysed whether the overlapping representation was useful in reducing the trial-to-trial variability in the image representation. Cortical neurons often show trial-to-trial variability in response to repetitions of the same stimulus. If neurons with spatially overlapping or similar representations showed independent or negatively correlated activity, the integration of activity among these neurons should reduce the variability in image representations across trials^36-38^.

The pattern of reconstructed image was reliably represented across trials regardless of the trial-to-trial response variability. Fig. 6a shows the trial-to-trial variability of the reconstructed images of the example case (shown in Fig. 5). Single-trial reconstructed images from all responsive neurons (55 cells) were generally stable across trials and were distorted in only a few trials (e.g., trial 10, Fig. 6a top). By contrast, the activity patterns of responsive neurons seemed variable across trials, although a few cells were reliably active (Fig. 6a bottom and 6b). To evaluate the trial-to-trial reliability of reconstructed images (or population responses), we used two measures, across-trial similarity and across-trial variability (see Methods). Across-trial similarity was defined as a Pearson’s correlation between single-trial reconstructed images (or response patterns) and their trial-average (Fig. 6c). Across-trial variability was a normalized squared error between single-trial reconstructed image (or response patterns) and their trial-average (Fig. 6d, see Methods). For all planes, the across-trial similarity of the reconstructed image pattern was high (0.85 [0.81–0.89], Fig. 6c left), whereas the across-trial similarity of the response was relatively low (0.40 [0.36–0.44], Fig. 6c right). Across-trial variability of the image was relatively low (Fig. 6d left), and that of the response pattern was relatively high (Fig. 6d right).

We next examined how the reconstructed images were reliably represented across trials. Based on the results of Fig. 5, neurons with partially overlapping or similar representation (reverse filters) will provide reliable representation across trials (for at least a part of the image), if these neurons are active on different trials. We demonstrate this using the example case (shown in Fig. 5). Among the nine overlapping cells, the three example neurons represented a partially similar pattern of the image (correlations of reverse filters among the three neurons were 0.31–0.68, Fig. 6e), while they were active on slightly different trials. Combining the nine overlapping cells resulted in the reliable representation of a local part of the image in most trials (Fig. 6e bottom). In the example case, the across-trial variability of a local part of the reconstructed image was gradually decreased as the number of cells used for the reconstruction increased (Fig. 6f). This tendency was also observed in data from all planes (Fig. 6g, black lines), suggesting that multiple overlapping cells support reliable image representation.

We further examined the relationship between reverse filter overlap or similarity and noise correlations (see Methods for a calculation, Supplementary Fig. 7a–c). Noise correlations were positively correlated with reverse filter correlations (correlation: 0.25, Supplementary fig. 7d) and also with signal correlations (correlation: 0.44, Supplementary fig. 7e)^37, 39^. However, noise correlations were tiny and mainly distributed near 0 even in cell pairs with high overlap or reverse filter similarity (e.g., in the pairs with correlation of reverse filter > 0.5, median noise correlation was 0.03. Supplementary fig. 7b and 7c). Furthermore, the maximal reverse filter similarity between a responsive cell and other overlapping cells was 0.57 [0.41–0.70] (Supplementary Fig. 6d), indicating that each responsive neuron often had at least one neuron with a similar reverse filter. Together, neurons with overlapping or similar reverse filters were almost independently active across trials, which was useful for reliable image representation.

**Figure 7.**
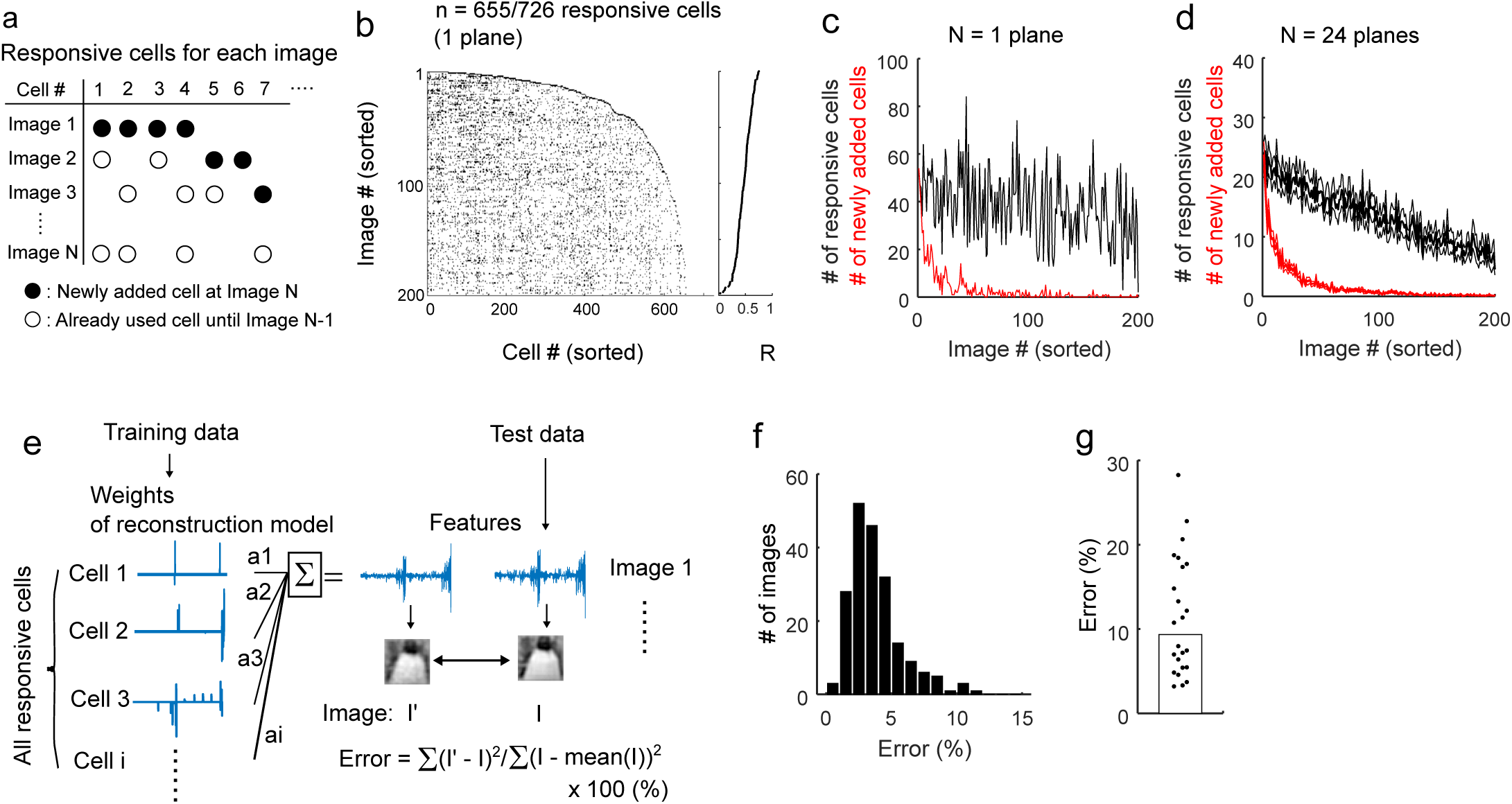
Visual features of natural images are distributed among most neurons in a population. **a**. Schematic of the analysis. Responsive neurons (open and closed circles) were plotted for each image. Closed circle: the responsive cells plotted for the first time at image N. Open circle: the responsive cells that have already plotted in image 1– (N-1). N: image number. **b**. Raster plots of highly responsive cells for each image in the representative plane shown in the previous figures (n = 655/726 responsive cells). The image # is sorted by the image reconstruction performance (descending order, right panel). In each line, cells that did not respond to the previously plotted images are added on the right side. As the image # increased, the number of newly added cells decreased, and then the cell # quickly reached a plateau level, indicating that many images are represented by a combination of cells that responded to other images. Thus, most images were represented with some degree of accuracy by a combination of subsets of responsive cells in the population. **c**. The numbers of responsive cells (black line) and numbers of newly added responsive cells (red line) are plotted against the image # for the case shown in (**b**). Again, the number of newly added cells quickly decreased as the image # increased. **d**. The numbers of responsive cells (black line) and numbers of newly added responsive cells (red line) are plotted against the image #. N = 24 planes. Three lines in each colour indicate the mean and the mean ± the standard errors. **e**. Schematic of the analysis. The feature set of each natural image was linearly regressed by the weights of the image reconstruction model (the cell selection model) from all the responsive cells in each plane. The weights of the reconstruction model were obtained based on a training dataset, and the target image was selected from a test dataset. The fitting error (% error, see Methods) was computed for each image on image space. If the features encoded in all responsive cells were sufficient to represent natural images, the weights of the responsive cells should work as basis functions to represent the visual features of the natural images. **f**. Distributions of the errors of all images in the example plane (shown in other figures). **g**. The median percent error (% error) across planes (bar, n = 24 planes). Each dot indicates the median of each plane.

We also analysed the effect of noise correlations on the reliable image representation across trials. Removing the noise correlations (see Methods) resulted in increases in the reconstructed performances and decreases in the across-trial variability (Supplementary fig. 7g–i), suggesting that the noise correlation is rather detrimental to both image reconstruction performances and the reliable image representation. In the relationship between the across-trial variability and the number of cells used for the reconstruction, the reduction of variability by removing the noise correlation was much smaller than that by increasing the cells used for the reconstruction (Fig. 6g), suggesting that noise correlation has a relatively small impact on the reliable representation.

### Representation of multiple natural images in a local population

Finally, we examined how multiple natural images were represented in a population of responsive neurons (Fig. 7a–7d). Figs. 7b and 7c show data obtained from the example plane presented in the previous figures (n = 726 cells). Natural images were sorted by reconstruction performance (y-axis in Fig. 7b and c), and the cells responding to each image are plotted in each row. First, as the number of images increased, new responsive cells were added, and the total number of responsive cells used for the reconstructions quickly increased (right end of the plot on each row, Fig. 7b). At approximately 50 images, the number of newly added responsive cells quickly decreased, and the increase in the total number of responsive cells slowed, indicating that the newly added image was represented by a combination of the already plotted responsive cells (i.e., the neurons that responded to other images), due to the small overlap in responsive cells between images (Fig. 1j). These findings are summarized in Figs. 7c and 7d, in which the number of newly added cells quickly decreased to zero as the number of images increased (red lines in Fig. 7c and 7d for the representative case and for all planes, respectively). Therefore, although only ∼5% of responsive neurons overlapped between images (Fig. 1j), this small overlap was useful for representation of many natural images by a limited number of responsive neurons.

We also analysed whether features encoded by the responsive neurons can represent any natural image as basis functions. If so, a set of features of the natural image will be accurately represented by a linear regression of the weights (i.e., features) of responsive cells in the cell-selection model independent of actual responses (see Methods, Fig. 7e). Fitting errors were computed in the image space. The median error was less than 10% for all images and all planes (3.3% [2.3–4.6%] for the example plane and 9.3% [5.4–18%], n = 24 for the planes, Fig. 7f and 7g). Based on this result, the features encoded by responsive neurons in a local population can accurately represent the visual contents of natural images.

### Image representation in awake mice

The results demonstrated thus far were obtained from anaesthetized mice. We finally examined whether the results were generalized to awake mice. We performed an additional experiment with awake, passively viewing mice that expressed GCaMP6s in the cortical cells (a transgenic mouse or mice introduced with GCaMP via virus vector, N = 7 planes from three mice)^40, 41^. We recorded eye positions and locomotion states (running and staying) during imaging and used data only when the eye was still (Fig. 8a–c, see Methods).

**Figure 8.**
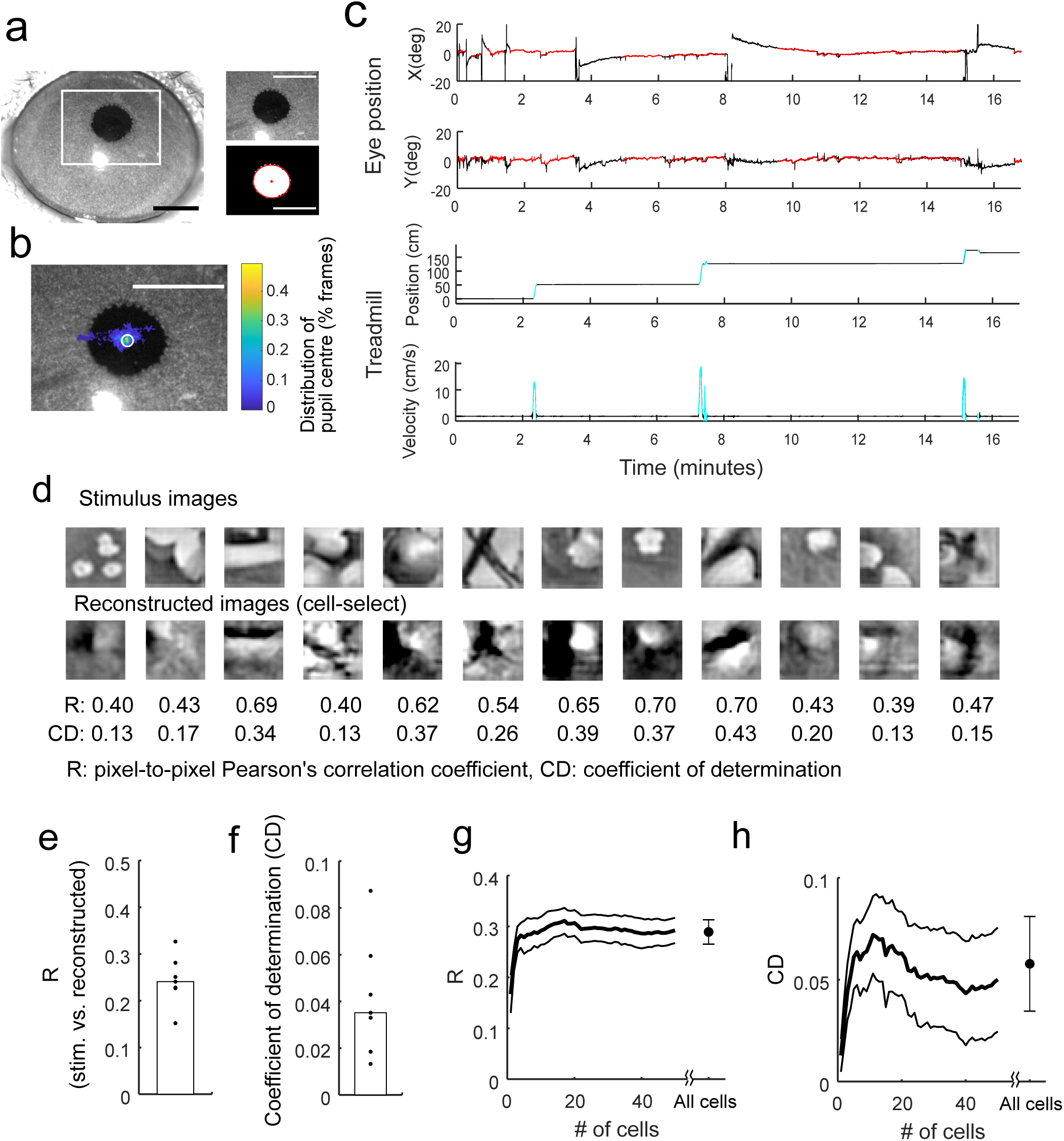
Image reconstruction in awake mice. **a**–**b**. Schematic of eye position analysis. (**a**) Image of a right eye. The white rectangle indicates an area recorded during imaging and analysed offline (left). The recorded image (upper right panel) was binarized, and the pupil was fitted with an ellipse (red contour in lower right panel). Centre of ellipse was used to estimate the eye position (red dot). Scale bars: 1mm. (**b**) Distribution map of eye position during imaging overlaid on the image in (**a**). The peak position of the distribution was detected, and data were used for subsequent analyses only when the eye stayed around the peak position (white circle, < 3.5 degrees or ∼70 microns on the image). Scale bar: 1 mm. **c**. Examples of eye position and locomotion state during imaging. Upper two panels: Horizontal (X) and vertical (Y) eye positions are shown. The red lines indicate time points at which eye stayed around the peak position (inside the white circle in **b**). Lower two panels: Position and velocity of a disc-type treadmill. The cyan lines indicate time points at which mice ran (velocity > 2 cm/sec). **d**–**h**. Image reconstruction by the cell-selection model in awake mice. (**d**) Examples of the reconstructed images. (**e** and **f**) Reconstruction performances (**e**: pixel-to-pixel correlation, R. **f**: coefficient of determination, CD). N = 7 planes. (**g** and **h**) R (**e**) and CD (**h**) against the number of neurons. A single image was reconstructed by a small number of neurons. The thick black line and grey lines indicate the means and the means ± standard errors, respectively (n = 6 planes). Data for images including at least 10 responsive cells were only used.

Results of main analyses obtained in the anaesthetized mice were also observed in the awake mice (Fig. 8d–h, Supplementary Figs. 8 and 9). Most cells were visually responsive for at least one image, whereas only a small fraction of cells were responsive for each image (Supplementary Fig. 8a–c). The percentages of responsive cells were 82% [78–94%] across stimuli (p<0.01 by ANOVA), and 1.5% [1.4–2.7] cells were responsive for each image (p<0.01 by t-test and the evoked response > 10%, Supplementary Fig. 8a and 8b). A single natural image was linearly decodable specifically from a small number of highly responsive cells for each image (Fig. 8d–h, Supplementary fig. 8d–i). Furthermore, reconstructed images were robust for cell drop (Supplementary fig. 8j and 8k) and reliable across trials (Supplementary Fig. 9). Therefore, the results obtained in anaesthetized mice were generalized at least to those in awake, passively viewing mice.

We also separately analysed the image reconstruction performance from excitatory and inhibitory cells. In some experiments, we used the mice that expressed tdTomato in inhibitory cells (gad2-cre × Ai14)^42, 43^ and identified excitatory and inhibitory cells based on the tdTomato (N = 5 planes from two mice). Excitatory cells were more visually responsive than inhibitory cells (Supplementary Fig. 10a–c). Furthermore, image reconstruction performances only by excitatory cells were comparable to those by all cells (Supplementary Fig. 10d and 10e). Therefore, excitatory cells mainly represent images at least under our experimental conditions.

We further examined how image reconstruction was affected by the locomotion state, which modulates V1 neuron activity^44, 45^. Although visual responses during running were higher than during staying (Supplementary Fig. 11a and 11b), the response patterns of a population of responsive cells were not largely different between the running and staying states (Supplementary fig. 11c). As a result, the reconstructed image patterns were not substantially affected by the locomotion states (Supplementary Fig. 11d and 11e).

## Discussion

In mouse V1, a single natural image activated a small number of neurons which was sparser than that predicted by the linear model. The Gabor features represented by the individual neurons only slightly overlapped between neurons, indicating diverse representations. The visual contents of natural images were linearly decodable mainly from a small number of highly active neurons, and the additional use of weakly active neurons degraded the decoding. The representation by a small number of neurons was achieved by their diverse representations. The image was reliably represented across trials regardless of trial-to-trial response variability likely due to multiple neurons with partially overlapping representation. Further, neurons with overlapping representation were almost independently active across trials, which should be beneficial to the reliable image representation. These image representations were observed in both anaesthetized and awake mice. Furthermore, a small overlap of responsive neurons between the images helped a limited number of responsive neurons to represent multiple natural images. Finally, the visual features represented by all responsive neurons provided a good representation of the original visual features in the natural images.

Visual responses to natural images or movies in V1 are sparse at the single cell level (high lifetime sparseness)^2, 3, 5-9^ and at the population level (population sparseness)^3, 5, 6, 14^. Recently, recordings of local population activity using two-photon Ca^2+^ imaging have enabled researchers to precisely evaluate the population sparseness^5, 14, 46^. We confirmed that a single natural image activated only a small number of neurons. According to the analysis of the encoding model, the visual responses of individual neurons were sparser than predicted from a linear model (Fig. 2f and 2g). Here, this sparse activity was shown to contain robust information to represent the natural image contents.

Image reconstruction is useful for evaluating the information content represented by neuronal activity and is widely used to analyse single unit activity in the LGN neurons^15^ and fMRI data from several visual cortical areas^16-19^ in response to natural scenes or movies. The former^15^ study used “pseudo-population” data collected from several experiments, and the latter studies^16-19^ used blood oxygen level-dependent (BOLD) signals that indirectly reflect the average local neuronal activity. However, it has not been examined whether and how the visual contents of natural images are represented in simultaneously recorded populations of single neurons in the cortex. It has also remained to be revealed how the information is optimally decoded from the representation; only a small number of highly active neurons mainly represent information, or the remaining neurons also have additional information. As shown in the present study, the visual contents of a single natural image were linearly decodable from a relatively small number of highly responsive neurons in a local population, and the additional use of the remaining neurons degraded the image reconstruction. Information has been proposed to be easily decoded from the sparse representation^4^. Indeed, the sparse population activity increases the discriminability of two natural scenes by rendering the representations of the two scenes separable^5^. Our results extend this finding by showing that information about visual contents represented in sparsely active neurons is linearly accessible, suggesting that downstream areas easily and optimally decode images by collecting activity of highly responsive neurons in the sparse representation of V1.

The visual features encoded by individual neurons should be diverse to ensure that a small number of active neurons represent the complex visual features of the image. Although RF structures in the local population of mouse V1 have already been reported^21, 22, 33, 34^, their diversity has not been analysed with respect to natural image representation. In the present study, the visual features represented by sparsely active neurons were sufficiently diverse to represent the visual contents of natural images (Fig. 7e–g). Computational models for natural image representation have suggested that sparse activity and the number of available neurons affect the diversity of RF structures^20, 47-49^.

Additionally, sparsely active neurons reliably represented an image across trials, regardless of trial-to-trial response variability. Although a computational model proposed sparse and overcomplete representation as the optimal representation of natural images by unreliable neurons^20^, this model has never been investigated experimentally. The reliable representation was mainly achieved by the diverse, partially overlapping representations, consistent with the overcomplete representation. Subregions of RFs of some V1 neurons partially overlap^21^. Based on our results, this overlap may be useful for reliable image representation. Furthermore, the reliable representation was also likely to be helped by neurons with similar reverse filters that were almost independently active across trials. Our results suggest a new representation scheme in which information is reliably represented while the representing neuronal patterns change across trials. This model seems to be similar to “drop-out” in deep learning^50^ and may be useful for avoiding overfitting and local minimum problems in learning.

Our analysis also revealed how multiple natural images were represented in a local population of responsive neurons. A single natural image activated specific subsets of neurons, whereas most neurons in a local population responded to at least one of the images, supporting the sparse, distributed code proposed in a previous study. Only 5.4% of responsive neurons overlapped between images (Fig. 1i). However, due to this small overlap, many natural images were represented by a limited number of responsive neurons (Fig. 7a–d). Furthermore, the features of all responsive neurons in a local population were sufficient to represent all the natural images used in the present study (Fig. 7e–g). Based on these findings, any natural image could be represented by a combination of responsive neurons in a local population. These findings also suggest that representation of multiple natural images is high dimensional, which is consistent with a recent report about high-dimensional representation in mouse V1^51^. Thus, a single natural image can be low-dimensionally represented in a high dimensional representation space for a large number of natural scenes.

In summary, this work highlighted how the visual contents of natural images are robustly represented in sparsely active V1 neurons. The diverse but partially overlapping representation helps the small number of neurons to robustly represent a complex image, regardless of across-trial variability. We propose a new representation scheme in which information is reliably represented with variable neuronal patterns across trials that may be effective in avoiding overfitting in learning, and that using the activity of only a small number of highly responsive cells is an optimal decoding strategy for the sparse representation.

## Methods

All experimental procedures were approved by the local Animal Use and Care Committee of Kyushu University and the University of Tokyo.

### Animal preparation for two-photon imaging in anaesthetized mice

C57BL/6 mice (male and female) were used (Japan SLC Inc., Shizuoka, Japan). Mice were anaesthetized with isoflurane (5% for induction, 1.5% for maintenance during surgery, ∼0.5% during imaging with a sedation of < 0.5mg/kg chlorprothixene, Sigma-Aldrich, St. Louis, MO, USA). The skin was removed from the head, and the skull over the cortex was exposed. A custom-made metal plate for head fixation was attached with dental cement (Super Bond, Sun Medical, Shiga, Japan), and a craniotomy (∼3mm in diameter) was performed over the primary visual cortex (centre position: 0–1 mm anterior to lambda, +2.5–3mm lateral to the midline). A mixture of 0.8 mM Oregon Green BAPTA1-AM (OGB1, Life Technologies, Grand Island, NY, USA) dissolved in 10% Pluronic (Life Technologies) and 0.025 mM sulforhodamine 101^52^ (SR101, Sigma-Aldrich) was pressure-injected with a Picospritzer III (Parker Hannifin, Cleveland, OH, USA) at a depth of 300–500 µm from the brain surface. The cranial window was sealed with a coverslip and dental cement. The imaging experiment began at least one hour after the OGB1 injection.

### Animal preparation for two-photon imaging in awake mice

In awake mouse experiments, we used two lines of transgenic mice: Thy1-GCaMP6s (GP4.3) transgenic mouse^41^ (JAX #024275, n = 1 mouse, 2 imaging planes) and the mice obtained by crossing gad2-ires-cre mice^43^ (JAX #010802) with Ai14 mice^42^ (JAX #007914) (Gad2-Ai14, n = 2 mice, 5 imaging planes). Thy1-GCaMP6s mice express GCaMP6s^40^ in cortical neurons. Gad2-Ai14 mice express tdTomato in almost all inhibitory neurons^43^.

GCaMP6s was introduced into Gad2-Ai14 mice via an adenoassociated virus (AAV). The Gad2-Ai14 mice were anaesthetized with isoflurane as described above. A small incision was made on a sculp, and a small hole (< 0.3 mm diameter) was made in the skull over V1. AAV2/1-syn-GCaMP6s^40^ (vector core; University of Pennsylvania, Philadelphia, PA, USA) was injected into V1 through the hole (titer: 3.0–5.0 × 10^12^ genomes/ml, volume: 500 nl, depth: 250-300 micron from a brain surface). After suturing the incision, the Gad2-Ai14 mice were recovered at least three days after the injection. The mice were anaesthetized with isoflurane to attach a metal plate for head fixation and to make a cranial window as described above. The mice were recovered after the surgery.

The mice were daily habituated with a head fixation on a disc type treadmill. Duration of the head fixation started with a few minutes and gradually prolonged up to ∼two hours over several days. If mice were calmly head-fixed for two hours without any stressful sign, imaging experiments started on the next day. The imaging started at least one week after the cranial window surgery, and three weeks after the virus injection for the gad2-Ai14 mice.

### Two-photon Ca^2+^ imaging

Imaging was performed with a two-photon microscope (A1R MP, Nikon, Tokyo, Japan) equipped with a 25× objective (NA 1.10, PlanApo, Nikon) and Ti:sapphire mode-locked laser (MaiTai Deep See, Spectra Physics, Santa Clara, CA, USA)^53, 54^. OGB1, SR101, GCaMP6s, and tdTomato were excited at a wavelength of 920 nm. Emission filters with a passband of 525/50nm were used for the OGB1 and GCaMP6s signals, and filters with a passband of 629/56nm for the SR101 and tdTomato signals. The fields of view (FOVs) were 338 × 338 µm (10 planes from 7 anaesthetized mice) and 507 × 507 µm (14 planes from 7 anaesthetized mice and 7 planes from 3 awake mice) at 512 × 512 pixels. The sampling frame rate was 30Hz using a resonant scanner.

### Monitoring of eye position and treadmill motion in awake mice

In awake mice experiments, right eye images and rotation of the disc type treadmill were recorded during the imaging. The right eye was monitored with a USB camera (NET New Electronic Technology GmbH, Germany). The treadmill rotation was monitored with a rotary encoder (OMRON, Japan). The eye images, the encoder signals and time stamps of frame acquisition of the two-photon imaging were simultaneously recorded using a custom-written program in LabView (National Instruments, Austin, TX, USA).

### Visual stimulation

Before beginning the recording session, the retinotopic position of the recorded FOV was determined using moving grating patches (lateral or upper directions, 99.9% contrast, 0.04 cycle/degrees, 2Hz temporal frequency, 20 and 50 degrees in diameter) while monitoring the changes in signals over the entire FOV. The lateral or upper motion directions of the grating were used to activate many cells because the preferred directions of mouse V1 neurons are slightly biased towards the cardinal directions^54, 55^. First, the grating patch of 50 degrees in diameter was presented in one of 15 (5 × 3) positions that covered the entire monitor to roughly determine the retinotopic position. Then, the patch of 20 degrees in diameter was presented on the 16 (4 × 4) positions covering an 80 × 80-degree space to finely identify the retinotopic position. The stimulus position that induced the maximum visual response of the entire FOV was set as the centre of the retinotopic position of the FOV.

A set of circular patches of greyscale, contrast-enhanced natural images (200 image types in main datasets, dataset 1) was used as the visual stimuli for predicting the response and reconstructing the natural image (256 intensity level, 60 degrees in diameter, 512 × 512 pixels, with a circular edge (5 degrees) that was gradually mixed to a grey background). Each natural image was adjusted to approximately full contrast (99.9%). The mean intensity across pixels in each image was adjusted to approximately 50%. Original natural images were obtained from the van Hateren Natural Image Dataset (http://pirsquared.org/research/#van-hateren-database)^56^ and the McGill Calibrated Colour Image Database (http://tabby.vision.mcgill.ca/html/welcome.html)^57^. Square image patches (512 × 512 pixels) were obtained from around centres of the original images. Some original images were down-sampled before the extraction of the centre parts. We selected 200 images that had spatial structure for the final stimulus set and did not include images that had less spatial structure (e.g., almost flat image) and very high spatial frequency components throughout the image (e.g., fine texture) by visual inspection. The pixel-to-pixel correlation between images was 0.003 [-0.12–0.11] (median [25–75^th^ percentile], n = 200 images).

During image presentation, one image type was consecutively flashed three times (three 200-ms presentations interleaved with 200 ms of grey screen), and the presentation of the next image was initiated after the presentation of the grey screen for 200 ms. Images were presented in a pseudo-random sequence in which each image was presented once every 200 image types. Each image was presented at least 12 times (i.e., 12 trials) for anesthetized and ∼40 times for awake mice in a total recording session. We did not set a long interval between image flashes to reduce the total recording time and increase the number of repetitions. In this design, the tail of the Ca^2+^ response to one image invaded the time window of the next image presentation (Fig. 1b). Although this overlap may have affected the visual responses to two adjacent images, many trial repetitions in the pseudo-random order and the sparse responses to natural images (Fig. 1) minimized the effects of response contamination between two consecutive images.

In another set of experiments with anaesthetized mice, we used different image sets (1000–2000 images that did not contain the 200 images described above, n = 4 planes from three mice). The original images for these image sets were obtained from the image datasets described above and from the Caltech 101 dataset (http://www.vision.caltech.edu/Image_Datasets/Caltech101/)^58^ and the web site for free image (https://www.pakutaso.com/). In experiments using these image sets, each image was presented 3–8 times.

Moving square gratings (8 directions, 0.04 cycles/degree, 2 Hz temporal frequency, 60-degree patch diameter) were presented at the same position as the natural image on the screen. Each direction was presented for 4 sec interleaved by 4 sec of the grey screen. The sequence of directions was pseudo-randomized, and each direction was presented 10 times for anaesthetized and 20 times for awake mice in a recording session.

All stimuli were presented with PsychoPy^59^ on a 32-inch, gamma-corrected LCD monitor (Samsung, Hwaseong, South Korea) with a 60-Hz refresh rate, and the timing of the stimulus presentation was synchronized with the timing of image acquisition using a transistor-transistor logic (TTL) pulse counter (USB-6501, National Instruments).

The entire recording session for one plane was divided into several recording sessions (4–6 trials/sub-session and 15–25 minutes for each session). Each recording session was interleaved by approximately 5–10 minutes of rest time, during which the slight drift of the FOV was manually corrected. In anaesthetized mice, the retinotopic position of the FOV was confirmed with the grating patch stimuli during the rest time every two or three sessions, and the recording was terminated if the retinotopic position had shifted (probably due to eye movement). The recordings were performed in one to three planes of different depths and/or positions in each anaesthetized mouse (1.7 ± 0.8 planes, mean ± standard deviation). In awake mice, the recording continued independent of the eye position and terminated if the mouse showed any stressful sign. The recording was performed in one plane per day, and one or two planes were obtained from each awake mouse.

For the analyses described below, natural images were scaled such that the maximum (255) and minimum (0) intensities were 1 and -1, respectively, and the grey intensity (127) was 0. A square (43 × 43 degrees) positioned in the centre of the natural image patch was extracted and down-sampled to a 32 × 32-pixel image. The down-sampled image was used to analyse the Gabor features, response prediction and image reconstruction.

### Analysis of eye positions and treadmill rotations

Each eye image in awake mice was binarized based on pixel intensities. The contour of the binarized pupil area was fitted with an ellipse whose centre was used as the eye position (Supplementary Fig. 7b). The eye positions on the image were transformed to angular positions. In the transformation, a previously reported value was used for the radius of the mouse eye^60^. In a distribution of the eye position during a total imaging session, a peak position was manually selected. Only the time points at which the eye was within 3.5 degrees (or ∼70 microns on the image) around the peak position were used for all analyses described below (except for the time course extraction of Ca signal from the two-photon imaging data).

In the analysis of the treadmill rotation, position signals from the rotary encoder on the treadmill were transformed to velocity and smoothed with the Savitzky–Golay filter. Running periods were defined as periods during which the velocity was greater than 2 cm/sec.

### Analysis of two photon imaging data

All data analysis procedures were performed using MATLAB (Mathworks, Natick, MA, USA). The recorded images were phase-corrected and aligned between frames. The average image across frames was used to determine the region of interest (ROIs) of individual cells. After removing the slow SF component (obtained with a Gaussian filter with a sigma of approximately five times the soma diameter), the frame-averaged image was subjected to a template matching method in which the two-dimensional, difference of Gaussians (sigma1: 0.26 × soma diameter that was adjusted for zero-crossing at the soma radius, sigma2: soma diameter) was used as a template for the cell body. Highly correlated areas between the frame-averaged image and the template were detected as ROIs for individual cells. ROIs were manually corrected via a visual inspection. SR101-positive cells (putative astrocytes^52^) were removed from the ROI in data of anaesthetized mice. For data of awake mice, a cross-correlation image^21^ and a max-projection image across frames were also used for the ROI detection.

The time course of the calcium signal in each cell was computed as an average of all pixels within an ROI. Signal contamination from an out-of-focus plane was removed using a previously reported method^54, 61^. Briefly, a signal from a ring-shaped area surrounding each ROI was multiplied by a factor (contamination ratio) and subtracted from the signal of each cell. In anaesthetized mice, the contamination ratio was determined to minimize the difference between the signals from a blood vessel and the surrounding ring-shaped region multiplied by the contamination ratio. The contamination ratios were computed for several blood vessels in the FOV, and the mean value for several blood vessels was used for all cells in the FOV. In awake mice, the contamination ratio was set to 0.7 for all cells following a previous study^40^, because it was difficult to identify the blood vessels in the GCaMP imaging.

After removing the out-of-focus signal, slow temporal frequency components (> 60 sec/cycle) were removed from the time course of each cell (a Gaussian low-cut filter applied on the frequency domain for anaesthetized data or a median low-cut filter for awake data), followed by smoothing with the Savitzky–Golay filter (4th order. 15 frame points length (∼500 ms)). Then, the filtered time course (F_filtered_) was transformed to ratio change (dF/F) by using the 20th percentile value across frames (F) (dF/F = F_filtered_ − F /F). Frame-averaged activity (dF/F) during 200 ms baseline (6 frames immediately before the stimulus) and during stimulus (average of the last 200 ms for each stimulus period) were used for subsequent analyses. The evoked response was obtained by subtracting the activity during baseline from that during stimulus. In awake mice, only data for images that contained at least six trials were used for subsequent analyses.

### Analysis of visual responses

Visually responsive neurons were determined by a one-way ANOVA (p < 0.01) with a dataset of N stimuli and one baseline (mean across stimuli) activity in each trial (size: N+1 activity × no. of trials, N: the number of stimuli used for the analysis). To validate this criterion, the ANOVA was applied to a randomized dataset in which data labels were shuffled in each trial. The false positive rate was only a small fraction of the percentage of the responsive cells (Anaesthetized: 85% responsive cells and 1.0% false positive rate. Awake: 82% responsive cells and 1.8% false positive rate).

For each responsive neuron identified by the ANOVA, responsiveness for each image was determined by using a t-test (p < 0.01, comparison of activity between stimulus and baseline) and a trial-averaged evoked response (> 10%). The evoked response threshold was used to reduce the false positive rate (Supplementary Fig. 1). The false positive rate was determined with the label-shuffled data. Without the evoked response threshold, the false positive rate was relatively high compared to the percentage of responsive cells per image (Anaesthetized: 3.2% for observed and 0.4% for shuffled data. Awake: 1.8% for observed and 0.3% for shuffled data). With the 10% amplitude threshold, the false positive rate decreased (Anaesthetized: 2.5% for observed and 0.1% for shuffled data. Awake: 1.5% for observed and 0.1% for shuffled data, Supplementary Fig. 1). Thus, we used the 10% evoked response threshold. We called the significant responses highly responsive. For the dataset with 1000–2000 stimulus images (dataset 2), responsive cells were not determined because of fewer trials.

The population sparseness (s) was computed using the equation described in previous studies^2, 3, 62^ as follows: s = [1 – (∑ Ri)^2^/ (N_cell_∑ Ri^2^)]/(1 − 1/N_cell_), where Ri is the evoked response of the ith cell, and N_cell_ is the number of cells (i = 1, …, N_cell_). Z-scored evoked responses were used in the following analyses including response prediction and image reconstruction (z-score was computed with responses across stimuli and trials in each cell).

### Gabor features

A set of spatially overlapping Gabor filter wavelets (n = 1248 filters) that has an almost self-inverting feature was prepared to extract the visual features of the natural images^10, 63, 64^. The down-sampled images were first subjected to the set of Gabor filters to obtain Gabor feature values. A single feature value corresponds to a single wavelet filter.

Gabor filters have four orientations (0, 45, 90, and 135 degrees), two phases, and 4 sizes (8 × 8, 16 × 16, 32 × 32, and 64 × 64 pixels) located on 11 × 11, 5 × 5, 3 × 3, and 1 × 1 grids (Fig. 2a and b). Therefore, the three smaller scale filters spatially overlapped with each other. The spatial frequencies of the four scale sizes of the Gabor wavelets were 0.02, 0.04, 0.09, and 0.18 cycle/degrees (cpd).

The Gabor filter set was almost self-inverting^63^, i.e., the feature values obtained by applying an image to the wavelet set were transformed to the image by summing the filters after multiplying by the feature values.

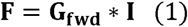

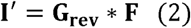

In equation 1 (eq. 1) **F** is the feature value matrix (matrix size: f × s. f: the number of features, 1248. s: the number of images), **G**_fwd_ is the Gabor filter matrix that transforms images to feature values and in which each row contains reshaped 2D-Gabor filters (f × p. p: the number of the image pixels, 1024), and **I** is the down-sampled stimulus image matrix (p × s). In eq. 2, **I**′ is the reconstructed image matrix (p × s) from **F, G**_rev_ is the Gabor filter matrix that transforms the features to images (p × f). In an ideal situation, **G**_fwd_^-1^ equals **G**_fwd_^T^ (i.e., **G**_rev_ = **G**_fwd_^T^. A^-1^ and A^T^ are inverse and transposed matrices of A, respectively.), resulting in that **I**^′^ equals **I**^63^. However, our Gabor transformation was not perfect; the pixel-to-pixel correlation between **I** and **I**’ was 0. 93 ± 0.026 (mean ± standard deviation, n = 200 images). To minimize the effect of this information loss on evaluations of image reconstruction performance (see below), we used **I**′ instead of **I** as target images for the evaluation of image reconstruction. The **G**_rev_ in eq. 2 was different from **G**_fwd_^T^ in eq. 1 in terms of a scaling and bias; **G**_rev_ = α × **G**_fwd_^T^ + β (α: scaling factor, β: bias. α and β were computed to minimize the sum of mean squared error between **I**′ and **I**). The Gabor filters and the transformations were based on an open source program (originally written by Dr. Daisuke Kato and Dr. Izumi Ohzawa, Osaka University, Japan, https://visiome.neuroinf.jp/modules/xoonips/detail.php?item_id=6894).

### Encoding model (response prediction model)

In the encoding model, single cell responses (**R**^k^ = [R_ki_], k: cell number, i: trial number across stimuli and trials. Size: 1 × N_trial_: the number of trials across stimuli and trials) were predicted using a linear regression analysis of selected Gabor feature values (**F**_select_ = [F_ji_], j: selected Gabor feature number. Size: f_select_ × N_trial_. f_select_: the number of selected features. Fig. 2a–c and Supplementary Fig. 2a and 2b).

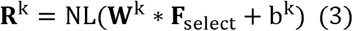

where **W**^k^ (= [W_kj_]. Size: 1 × f_select_) is the weight vector, **F**_select_ (f_select_ × N_trial_) is the matrix of selected feature values, b^k^ (Size: 1 × 1) is bias, and NL( ) is the non-linear scaling function. The encoding model was created independently for each cell. The features used in the regression were determined as follows. First, Pearson’s correlation coefficients between the response and feature values were computed in each feature. Then, using one of the preset values for the correlation coefficient as a threshold (13 points ranging from 0.05 to 0.35, Supplementary Fig. 2a and 2b), only the more strongly correlated features were selected (feature selection) and used in the regression analysis. **W**^k^ and b^k^ were estimated to minimize the loss function: 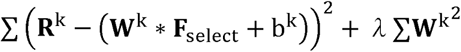: (*λ*: regularization parameter). This was solved by using Bayesian linear regression with an expectation-maximization algorithm that is approximately equivalent to linear regression with L2 regularization^65^. After the regression analysis, the non-linearity of the predicted response was adjusted with a rectification step using the following function^34^, NL(x): NL (x) = A/[1 exp (Bx C)] D, where A, B, C and D are parameters to be estimated using a built-in Matalb function (*lsqnonlin*). This step merely scaled the regression output without changing the regression parameters (**W**^k^ and b^k^).

The response prediction performance of the model was estimated by 10-fold cross-validations (CVs) in which the response data for 90% images were used to estimate the parameters, and the remaining data for 10% images were used to evaluate the prediction (Thus, **W**^k^ and b^k^ were estimated and fixed in each CV). In the 10-fold CVs, all images were used once as test data. The prediction performances were estimated using Pearson’s correlation coefficients between the observed (trial-average) and predicted responses. Encoding models were created for all preset threshold values for feature selection, and the model that exhibited the best prediction performance was selected as the final model.

In the analysis of overlapping weights (i.e., feature) between two cells, the percentage of overlapping weights relative to the number of non-zero weights was computed for each cell and averaged between the two cells in the pair.

Using the same dataset as used in the encoding model, the RF structure was estimated for each cell using a regularized inverse method^32-34^ that employs one hyper-parameter (regularized parameter). In the 10-fold CVs, the RF structure was estimated with the training dataset using one of the preset regularized parameters (13 logarithmically spaced points between 10^−3^ and 10^3^). The visual response was predicted using the estimated RF and test dataset. The prediction performance of visual response was estimated by determining Pearson’s correlation coefficients between the observed and the predicted responses. RFs were estimated for all values of the preset regularized parameters, and the value that resulted in the best predicted response was selected for the final RF model.

### Image reconstruction

For image reconstruction, feature values obtained from each Gabor filter were linearly regressed by the single-trial activity of multiple cells. For each Gabor feature,

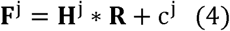

where **F**^**j**^ (= [F_ji_]) is the feature values for jth Gabor filter (j: Gabor feature number, i: trial number across stimuli and trials. Size: 1 × N_trial_. N_trial_: the number of trials across stimuli and trials), **H**^**j**^ (= [H_jk_]. Size: 1 × N_cell._ N_cell_: the number of cells) is the weights, **R** (= [R_ki_]. k: cell number. Size: N_cell_ × N_trial_). is the response matrix. In the 10-fold CVs, the weights, **H**^j^ and a bias, c^j^ were estimated to minimize the loss function: 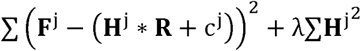, which was solved by using Bayesian linear regression with an expectation-maximization algorithm with the training dataset. Then, each Gabor feature value 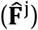 was reconstructed from the visual responses in the test dataset (10-fold CV with the same data split to that in the encoding model. **H**^**j**^ and c^j^ were estimated and fixed in each CV). After each Gabor feature was independently reconstructed, sets of reconstructed feature values (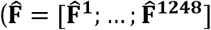. Size: f × N_trial_) were transformed into images using eq. 2.

In the all-cell model, each feature was reconstructed using all cells (Fig. 3a, left panel). In the cell-selection model, each feature was reconstructed using a subset of cells. For each feature reconstruction, cells were selected using the encoding model; if a cell was represented by jth feature in the encoding model (i.e., non-zero weight in jth feature in the eq. 3), the cell was selected for the jth feature reconstruction (Fig. 3a, right). In other words, each cell participated in the reconstruction of features that the cell encoded. When none of the cells were selected for feature reconstruction, the feature value was set to 0.

Reconstruction performance was evaluated using pixel-to-pixel Pearson’s correlation coefficients (R) and coefficients of determination (CD) between the stimulus and reconstructed images. CD was computed using the following equation: CD = 1 − ∑(**I**′ − **Î**)^2^/∑(**I**′ − **I**′_mean_)^2^ (**Î** the reconstructed image by the model, **I**′: stimulus image obtained by the transformation and reconstruction of Gabor filters (eq. 2), **I**′_mean_: mean pixel intensity of **I’**). R indicates similarity of image pattern between **Î** and **I**′, and CD indicates goodness of model prediction reflecting differences in pixel intensities between **Î** and **I**′.

The cell-selection described above (i.e., feature selection in the encoding model) should overestimate the reconstruction performance, because the test dataset was used for both the cell-selection and the performance evaluation of the reconstruction model. To precisely evaluate the performance of the cell-selection model, we used nested CV for the cell-selection; a dataset was separated into 10% test, 9% validation, and 81% training sets, and the cell-selection was performed with the validation and training sets. Then, the performance of the reconstruction model that was trained with both the validation and training sets was evaluated using the test dataset. The performance of the reconstruction model with nested CV was similar to that of the model without nested CV (Supplementary Fig. 4).

In the analysis of the overlapping weights (i.e., feature) between cells, the percentage of overlapping weights relative to the number of non-zero weights was computed for each cell and averaged between the two cells in the pair.

We independently obtained the weights of the image reconstruction model (**H** = [**H**^1^; … ; **H**^1248^], size: f × N_cell_), the weights of encoding model (**W** = [**W**^1^; … ; **W**^Ncell^]^T^, size: f × N_cell_) and RF by the pseudoinverse method. We chose the scheme of the image reconstruction model to optimally reconstruct the image by a population of neurons. In the image reconstruction model, **H** was estimated to directly optimize the image reconstruction considering the responses of multiple cells. By contrast, in the encoding model (or RF estimation), weights were estimated independently in each cell without considering the other cells’ responses. Because **H** was likely to be more optimized to represent images with multiple cells than **W** or RF, we chose the model scheme for the image reconstruction.

### Relationship between the image reconstruction and the number of features that each neuron encoded

In the analysis in Supplementary Figs. 5a–c, the cell-selection model was used for the image reconstruction. In the cell-selection model, each neuron participated in the reconstructions of a small number of features that were strongly correlated with the neuron’s responses. In the cell-selection model shown in Fig. 3 (original model), the threshold for the correlation coefficient was selected based on the encoding model for each neuron (Fig. 3a, right panel). In each neuron, the threshold of correlation coefficient was adjusted to increase (or decrease) the number of features for which each neuron participated in their reconstruction (0.1–20 fold change in the number of features per neuron relative to the original model, Supplementary Fig. 5a–c). In each fold change, the reconstruction model was trained with training data (i.e., weights and bias parameters were estimated in each fold change of the number of features), and the performance was estimated with test data using the 10-fold CV as described above.

### Image reconstruction from a small number of cells

In the analyses shown in Figs. 4a–4h and 5a–5b, cells in each image were separated into responsive and remaining cells and sorted by their response amplitude in descending order (i.e., from highest to lowest response amplitude). Then, the cells were selected first from the responsive cells and then from the remaining cells for the addition (Fig. 4a–h) or dropping (Fig. 5a–b). The analyses only used data for images including at least 10 responsive cells in Fig. 4a–h and at least 5 responsive cells in Fig. 5a–d.

In the image reconstruction from a subset of cells for each image (Fig. 4, 5 and 6), the weights of the cell-selection model (**H** = [**H**^1^; … ; **H**^1248^], f × N_cell_) were scaled because **H** was estimated by the more cells than the cells used during the cell addition or dropping.

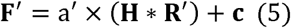

where **F**′ (f × N_trial_) is the matrix of all reconstructed feature values, **H** and **c** ([= [c^1^;… ; c^1248^], f × 1) are the weight and bias matrices of the cell-selection model in eq. 4, **R**′ is a response matrix that includes a subset of cells used for each image reconstruction (i.e., the responses of non-selected cells were set to 0), and a′ is a free parameters that is obtained to minimize the sum of squared error between the original and reconstructed feature values across all features and stimuli of the training dataset in each CV: ∑(F′ –F)^2^, (**F**: a matrix of features of regression target). Because a′ is common across all features, this scaling did not change the weight pattern of the cell-selection model. Then, images were reconstructed from F′ using eq. 2 as described above. In the reconstruction from a subset of cells (Fig.4, 5, and 6), a′ (i.e., weights, a′ × **H**) was estimated independently for each subset of cells, and a different set of cells was used for each image.

### Robustness of image reconstruction against cell drop

In the analysis of robustness (Fig. 5c–5f), a representation area of each cell was determined using the z-scored reverse filter (sum of weights × Gabor filters). The representation area was defined as a cluster of pixels whose absolute z-scores were greater than 1.5 and whose contours were smoothed (e.g., red contours in Fig. 5c and Supplementary Fig. 6a). If multiple areas were obtained, the largest was used. Then, using the representation area, the overlap index was computed between responsive cells in each stimulus; overlap index = (A∩B)/(A∪B), where (A∩B) is the overlapping representation area between cell A and cell B, and (A∪B) is a combined representation area between cell A and cell B (Supplementary Fig. 6a). Using the overlap index, a set of overlapping cells was selected for each responsive cell; the overlapping cells consisted of one responsive cell (reference cell) and other responsive cells that overlapped with the reference cell (overlap index > 0.2). This analysis did not care whether other overlapping cells overlap each other or with other non-selected cells.

To evaluate the effects of cell drop, cells were randomly removed from the overlapping cells, and the reconstructed image was computed after each cell was dropped. The reference cell was initially removed, and then other remaining overlapping cells were removed in each cell drop sequence. Changes in the reconstructed images were estimated by quantifying the pixel-to-pixel correlation (R) of a local part of the image. The local part of the image was determined as the representation area of the reference cell that was overlapped by the area of at least one of other overlapping cells (overlapping area in Fig. 5c). A random drop of overlapping cells was repeated 120 times, and the results were averaged across the random orders in each reference cell. All responsive cells were used once as the reference cell in each stimulus image. This analysis only used data for images including at least 5 responsive cells and sets of overlapping cells including at least 5 overlapping cells.

### Across-trial similarity and variability

To estimate the reliability of reconstructed image (or response patterns) across trials, two measures were used: across-trial similarity and across-trial variability. For the across-trial similarity of reconstructed images (or response patterns) (Fig. 6c), Pearson’s correlations between single-trial reconstructed images (or response patterns) and their trial-average were computed and averaged across trials.

For the across-trial variability (Fig. 6d–g), normalised squared errors between single-trial images (or response patterns) and trial-averaged images (or response patterns) were computed using the following equation: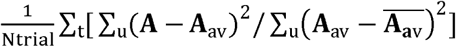 (**A**: single-trial reconstructed image or response patterns,**A**_av_: trial-averaged reconstructed image or response pattern, 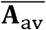: mean of **A**_av_ across pixels or cells. Ntrial: the number of trials. ∑_t_: summation across trials. ∑_u_ : summation across pixels or cells).

In the analysis in Fig. 6f and 6g, the overlapping cells were selected as described above for each responsive cell (i.e., the reference cell. See the section of **Robustness of image reconstruction against cell drop**). The reference cell was initially selected, and then other overlapping cells were randomly selected for a set of cells that used for image reconstruction (the sequences of random cell selection was repeated 200 times.). The image was reconstructed from the subset of overlapping cells, and across-trial variability of a local part (i.e., overlapping area) of the reconstructed image was computed for each subset of cells. Only data for images that contained at least 5 responsive cells were used in the analyses in Fig. 6.

### Noise correlation

In the analysis in Supplementary Fig. 7, the noise correlation was computed using responses across stimuli. Evoked responses in each stimulus image were transformed to z-score and collected across stimuli in each cell. Then, Pearson’s correlation coefficient was computed between the collected responses in a cell pair and used as the noise correlation. To remove the noise correlation, responses to each stimulus were shuffled across trials independently in each cell. Using the shuffled data, image reconstruction model was obtained as described above for the analyses in Fig. 6g, Supplementary Fig. 7g–i, 7p–r, and Supplementary Fig. 9f.

### Capacity of image representation by encoded features in a population

The analysis in Fig. 7e–g examined whether the features encoded by responsive cells could represent image as a basis function independent of actual neural responses. If the features encoded by responsive neurons can represent any image, a set of features of a given image (**F**) will be linearly regressed by the weights of responsive cells in the cell-selection model, **H** (= [**H**^1^; … ; **H**^1248^], f × N_cel_. **H**^k^ in eq. 4),

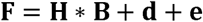

where **B** and **d** are free parameters that are calculated to minimize the sum of squared error, ∑(**F**– (**H** * **B** + **d**))*2*, and **e** is an error term. **F** was selected from a test dataset and **H** was obtained from a training dataset in the 10-fold CV. The fitting was evaluated by calculating the fitting error (error) on the image space as follows:

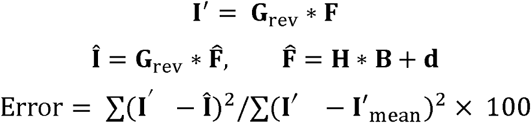

where **G**_rev_ is the Gabor filter matrix for reconstruction (eq. 2), and **I**’mean is the mean pixel intensity of **I**’. Thus, this analysis estimated how well the features of individual neurons in a local population could represent the image features independent of actual neuronal activity. In other words, this analysis estimated the upper bound capacity of a local population to represent any image with an ideal combination of cell features (with parameters B and **d**).

### Effects of locomotion state on image reconstruction

Because the awake mice were not trained to run, they often stayed calmly during imaging. In the analyses shown in Supplementary Fig, 11, we included only data from two planes in one mouse (Thy1-GCaMP6s mouse) that ran relatively frequently. Furthermore, we used only data for images that contained at least five responsive cells, four running trials and four staying trials (80 image cases, n = 295 responsive cells). The running modulation index (RMI) for each cell was defined as follows: RMI = (R_run_-R_stay_)/(R_run_+R_stay_), where R_run_ and R_stay_ were the mean evoked responses during running and stay periods, respectively. RMI was computed in each responded image and averaged across images in each cell. Image reconstruction was performed using data with both conditions, and the performances were collected separately in each condition (Supplementary Fig. 11d and e).

## Statistical analyses

All data are presented as the median and 25–75^th^ percentiles unless indicated otherwise. The significant level was set to 0.05, with the exception of the criteria of significant visual response (0.01). When more than two groups were compared, the significant level was adjusted with the Bonferroni correction except for the visually responsive cell analysis. Two-sided test was used in all analyses. The experiments were not performed in a blind manner. The sample sizes were not predetermined by any statistical methods but are comparable to the sample size of other reports in the field.

## Data availability

The datasets of the current study and the code used to analyse them are available from the corresponding authors on reasonable request.

## Acknowledgements

We thank Ms. Y. Sono, A. Hayashi, T. Inoue, A. Ohmori, A. Honda, M. Nakamichi for animal care, and all members of Ohki laboratory for support and discussions. This work was supported by grants from Core Research for Evolutionary Science and Technology (CREST)—Japan Agency for Medical Research and Development (AMED) (to K.O.), Japan Society for the Promotion of Science (JSPS) KAKENHI (Grant number 19H05642, 25221001 and 25117004 to K.O. and 15K16573, 17K13276 to T.Y.), Brain Mapping by Integrated Neurotechnologies for Disease Studies (Brain/MINDS)—AMED (to K.O.), Strategic International Research Cooperative Program (SICP)—AMED (to K.O.), grants from the Ichiro Kanehara Foundation for the Promotion of Medical Sciences and Medical Care, and the Uehara Memorial Foundation (to T.Y.).

## Author contributions

T.Y. and K.O. designed the research. T.Y. performed experiments. T.Y. and K.O. analysed data and wrote the manuscript. K.O. supervised the research.

