## Supplementary Information for "Robust representation of natural images by sparse and variable population of active neurons in visual cortex"

### Contents

Supplementary Figure 1.

Supplementary Figure 2.

Supplementary Figure 3.

Supplementary Figure 4.

Supplementary Figure 5.

Supplementary Figure 6.

Supplementary Figure 7.

Supplementary Figure 8.

Supplementary Figure 9.

Supplementary Figure 10.

Supplementary Figure 11.

**Supplementary Figure 1. Fraction of responsive cells for each image and false positive rates with three levels of amplitude thresholds**

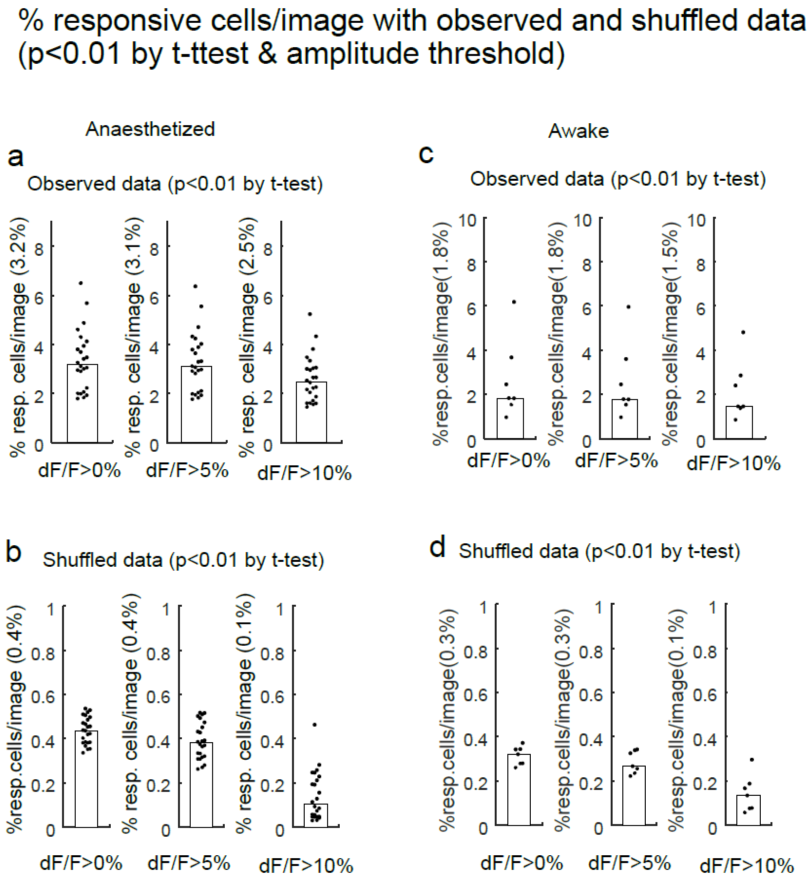


**Supplementary Figure 1. Fraction of responsive cells for each image and false positive rates with three levels of amplitude thresholds**

**a** and **b.** Percentages of responsive cells per image in anaesthetized mice. In each responsive neuron identified using ANOVA ($p<0.01$), responsiveness for each image was determined using a t-test ($p<0.01$, baseline vs. stimulus activity) and evoked response amplitudes. **a**. The percentages of responsive cells per image obtained from observed data with three levels of the evoked response thresholds (0%, 5%, and 10%). **b**. False positive rate estimated with label-shuffled data. The 10% threshold resulted in a small fraction of the false positive rate relative to the responsive rate obtained from the original data.
**c** and **d**. Percentages of responsive cells per image in awake mice. Same as in (a) and (b), except for awake data

**Supplementary Figure 2. Gabor filter set and prediction performance of encoding model**

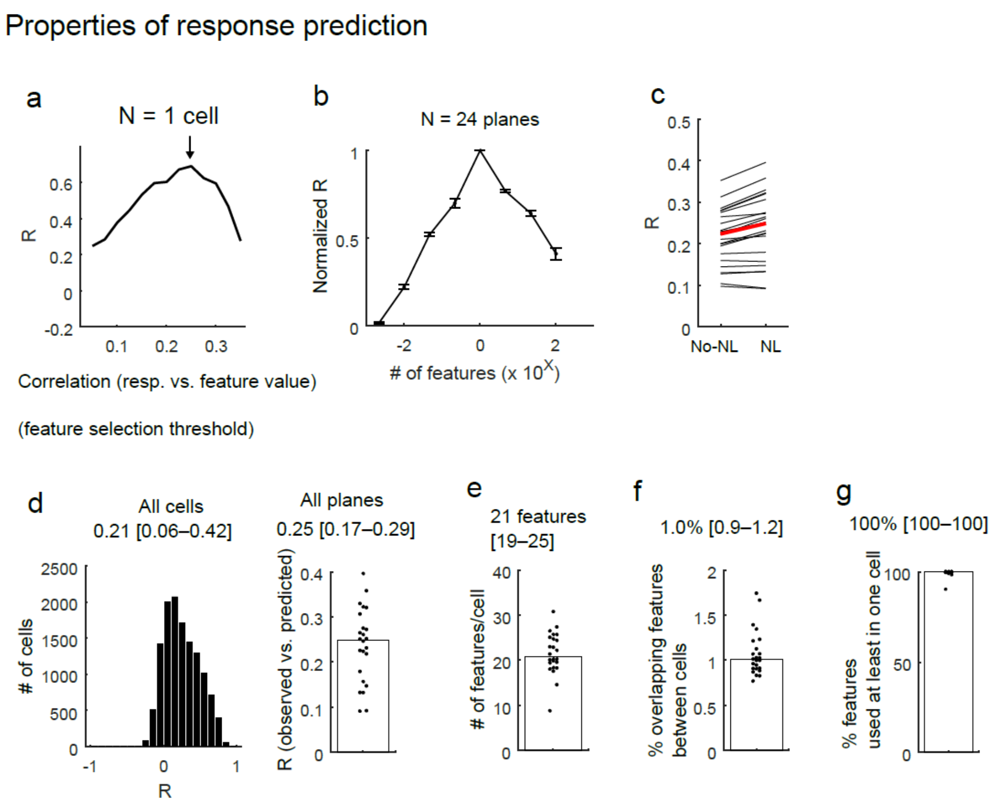


**Supplementary Figure 2. Gabor filter set and prediction performance of encoding model**

**a**. Effect of feature selection on the response prediction performance of the encoding model in an example cell. The response prediction performances (correlation coefficients between the observed and predicted responses, R) are plotted against the threshold values of the feature selection. First, Pearson's correlation coefficient between each feature value and response was computed. Then, the features with correlation coefficients greater than the preset threshold values (x-axis) were used for the regression analysis of the encoding model. The threshold value for the final model was selected from the values to maximize the response prediction in each neuron (arrow).
**b**. Normalized prediction performance against the number of features used for the encoding model. The data plotted in (**a**) were normalized by the maximal performance (for the y axis) and re-plotted against the number of features used in the model of each threshold (for the x-axis). Data were collected across all cells in each plane, re-sampled and averaged in each bin (x-axis in the plot). The plotted curve was averaged across planes. Means ± standard errors are shown (n = 24 planes).
**c**. Response prediction performances of the encoding models with and without a non-linear scaling step. R: correlation coefficient between observed and predicted responses. No-NL: model without a non-linear scaling step. NL: model with a non-linear scaling step. No-NL: 0.22 [0.17–0.27]. NL: 0.25 [0.17–0.29]. P = $1.1 \times 10^{-4}$ by signed-rank test (n = 24 planes).
**d**. Distributions of the response prediction performances for all cells (0.21 [0.06–0.42], n = 12,755 cells, left panel) and for all planes (0.25 [0.17–0.29], n = 24 planes, right panel).
**e**. The number of features encoded by each cell (21 [19–25] features, n = 24 planes).
**f**. Percentages of overlapping features between cells (1.0% [0.9–1.2%], n= 24 planes). In each cell pair, the number of overlapping features was divided by the number of features used in the encoding model for each cell and averaged between the two cells in a pair.
**g**. Percentages of features that were used for response prediction of at least one cell in a population (100% [100–100%], n = 24 planes).

**Supplementary Figure 3. Properties of Gabor features encoded by individual cells.**

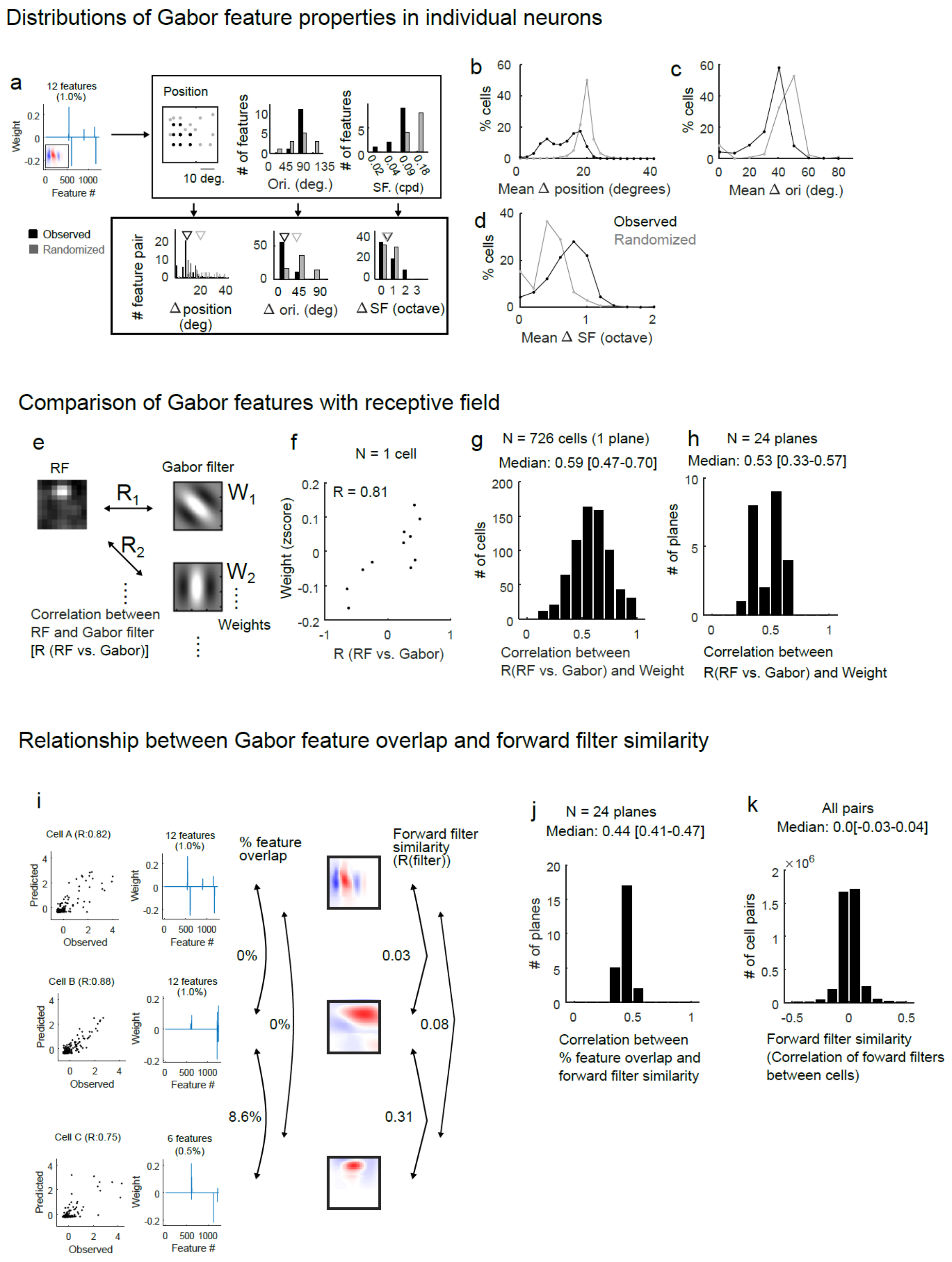

**Supplementary Figure 3. Properties of Gabor features encoded by individual cells.**

**a–d.** Properties of Gabor features encoded by individual cells.
**a**. Schematic of the analysis. Upper panels: Features encoded by each cell were analysed in terms of retinotopy (position, **b**), orientation (Ori, **c**) and spatial frequency (SF, **d**). Lower panels: similarities among features in each cell were evaluated by computing differences in positions, orientation, and spatial frequency between features. Mean values in each cell were collected across all cells and plotted in (**b–d**). As a control, labels of Gabor features were randomized in each cell while keeping the number of features (randomized).
**b–d**. Distribution of mean differences in position (**b**), orientations (**c**), and spatial frequency (**d**). Black and grey lines indicate the results of observed and randomized data, respectively. Relatively narrow distributions of randomized data reflect bias of original Gabor filter properties (e.g., slightly more Gabor filters are located at the centre positions, see Supplementary Fig. 2b).

**e–h**. Relationship between the weights of the Gabor features and the RF structure.
**e** and **f**. Schematics of the analysis. (**e**) In each cell, the RF structure was determined using a pseudo-inverse method (see Methods), and pixel-to-pixel Pearson's correlation coefficients between the RFs and Gabor filters were computed (R1 and R2 in (**e**); R (RF vs. Gabor)). (**f**) Then, Pearson's correlation coefficient between the R (RF vs. Gabor) and weight values (W1 and W2 in (**e**)) was computed for each cell. In the example cell shown in (**f**), the R (RF vs. Gabor) and weight values were positively correlated (correlation coefficient: 0.81), indicating that high weight value were assigned to Gabor filters similar to the RF.
**g** and **h**. Distributions of the correlation coefficient between R (RF vs. Gabor) and weights. Distributions for all cells in the example plane (**g**) and for all planes (**h**).

**i–k**. Relationship between the percentage of overlapping features and the encoding filter similarity in a cell pair.
**i**. Examples of response prediction performances (left panels), weights (centre) and encoding filters (right panels) of the three cells (cell A, B and C). Among the three cells, the percentages of overlapping non-zero features (% overlapping feature) were 0%, 8.6%, and 0% (for cell pairs A-B, B-C and A-C), whereas the similarities of the forward filters (pixel-to-pixel correlation) were 0.03, 0.31, and 0.08 for the three pairs. The encoding filter was obtained by computing the sum of Gabor filters multiplied by the weights of the encoding model. The red and blue colours of the encoding filter indicate positive and negative values, respectively. The correlation coefficient between the % overlapping features and encoding filter similarity was computed in each plane and plotted in (**j**).
**j**. Distribution of the correlation coefficient between the % overlapping feature and the absolute value of encoding filter similarity (n = 24 planes).
**k**. Diverse structure of the encoding filters. The similarity of the encoding filter was relatively small for all pairs (median: 0.0), which indicated the diverse structure of the encoding filters between cells.

**Supplementary Figure 4. Comparison of image reconstruction performances between the models with and without nested CV**

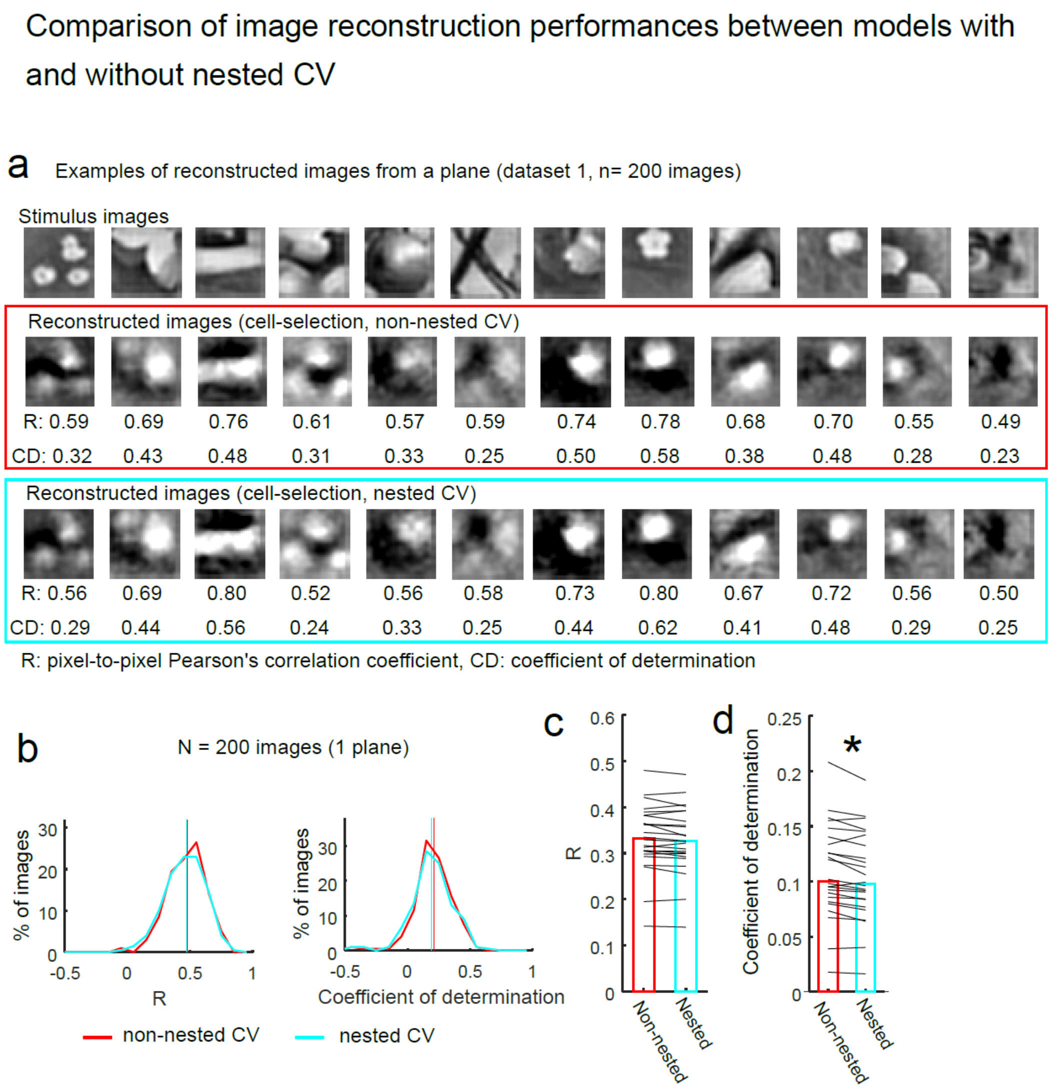


**Supplementary Figure 4. Comparison of image reconstruction performances between the models with and without nested CV**

**a**. Examples of reconstructed images from main datasets (dataset 1, 200 images). Stimulus images (top), images that were reconstructed using the cell-selection model without nested CV (non-nested CV, middle, also shown in Fig. 3c in the main text) and using the cell-selection model with nested CV (nested CV, bottom) are shown. Each reconstructed image was averaged across trials. The reconstruction performances (R and coefficient of determination, CD) were computed for each trial, and trial-averaged performances are presented below each reconstructed image.
**b**. Distributions of R (left) and CD values (right) for the model without nested CV (red lines) and the model with nested-CV (cyan lines) in the example plane shown in Figs. 1 and 2 (n = 200 images reconstructed using 726 cells from a plane). Vertical lines indicate median values.
**c** and **d**. R (**c**) and CD (**d**) of dataset 1 across planes. *: p = 0.001 in (**d**) using the signed-rank test (N = 24 planes). The reconstruction performances of the cell-selection model with nested CV were similar to those with non-nested CV.

**Supplementary Figure 5. Reconstruction performances against the number of features per cell and spatial frequency components**

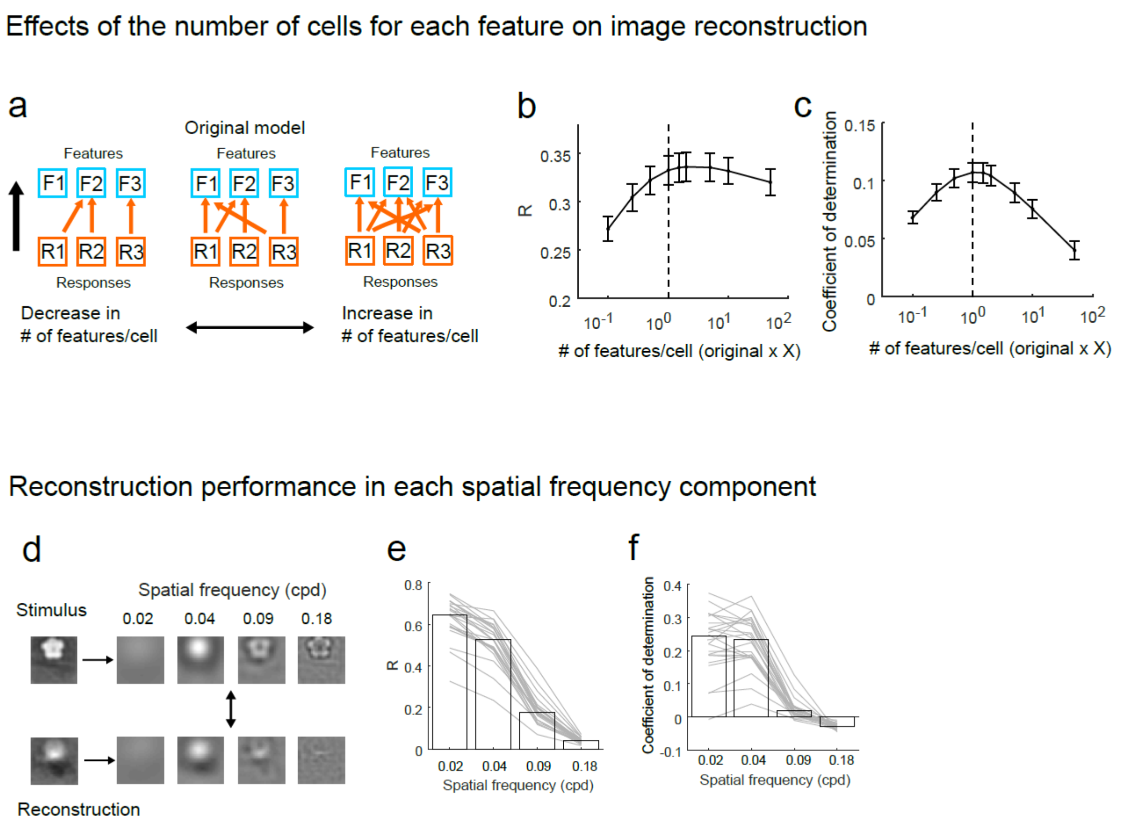


**Supplementary Figure 5. Reconstruction performances against the number of features per cell and spatial frequency components**

**a.** Schematic of the analyses. This analysis examined how the image reconstruction performance was affected by the change in the number of features of which each cell participated in the reconstruction.
**b** and **c.** Reconstruction performances (R in **b,** and coefficient of determination in **c**) against the number of features per cell. The number of features of the original model (shown in the main text) was indicated by the dotted vertical line (x = original × 1) in each panel. The original model demonstrated nearly optimal performance, suggesting that how the individual neurons encode Gabor features is one of the important factors of image reconstruction and that the original model captures nearly optimal feature-cell assignment.
**d–f.** Reconstruction performance in each spatial frequency component by the cell-selection model. (**d**) Schematic of the analysis. Images were reconstructed using features of each spatial frequency. (**e** and **f**) Pixel-to-pixel correlation coefficient (R, **e**) and coefficient of determination (**f**) were plotted (n = 24 planes). Relatively low spatial frequency components of images were represented by V1 neurons.

**Supplementary Figure 6. Spatial overlap of reverse filters among responsive neurons**

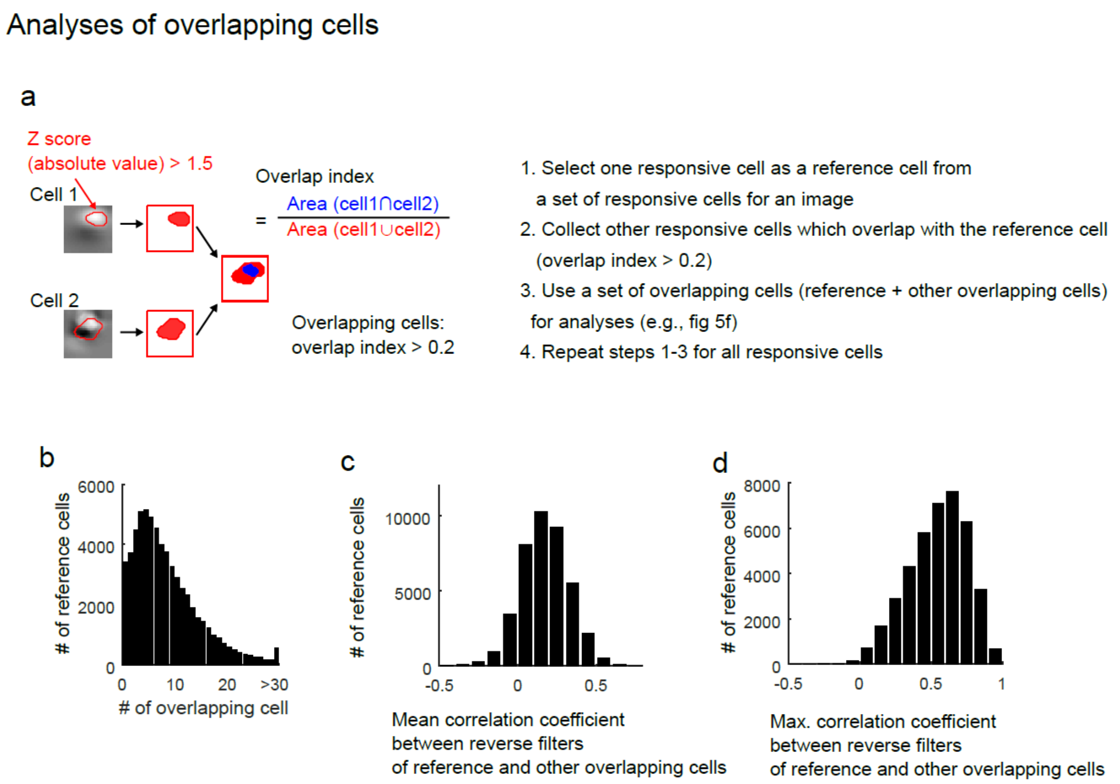


**Supplementary Figure 6. Spatial overlap of reverse filters among responsive neurons**

**a**. Spatial overlap of reverse filters. The reverse filter was transformed to a z-score, and the area in which the absolute z-score was greater than 1.5 was defined as a significant area (red contours in the left panels). The significant area was used to compute the overlap index. For each responsive cell (reference cell), the overlap index between the reference cell and other responsive cells was computed, and other responsive cells whose overlap indices were greater than 0.2 were collected for a set of overlapping cells (reference and the collected cells).
**b**. Distribution of the number of spatially overlapping cells for each responsive cell.
**c**. Distribution of the mean correlation coefficient between reverse filters of a reference cell and other overlapping cells.
**d**. Distribution of maximal correlation coefficient between reverse filters of a reference cell and other overlapping cells.

**Supplementary Figure 7. Relationship between noise correlation and reverse filter similarity**

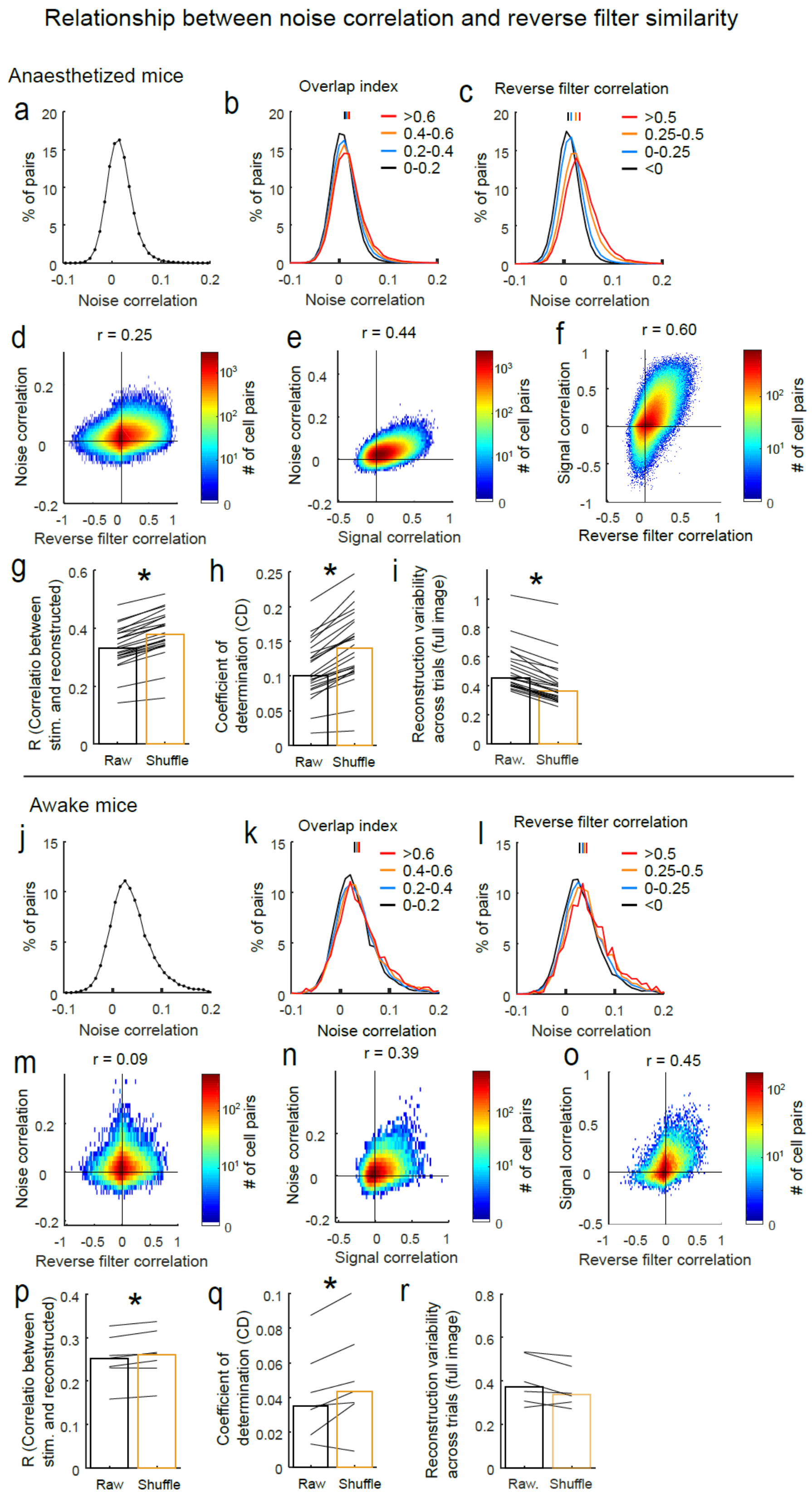

**Supplementary Figure 7. Relationship between noise correlation and reverse filter similarity**

**a**. Distribution of noise correlation from all responsive cell pairs (0.014 [-0.002–0.031]) in anaesthetized mice.
**b** and **c**. Distributions of noise correlations grouped by overlap index (**b**) and reverse filter similarity (pixel-to-pixel correlation of reverse filters, **c**). The distribution in (**a**) was divided into four groups based on the overlap index or the reverse filter correlations, and the distributions of the groups were plotted with different colours. Vertical lines over distribution curves indicate median values. In the analyses in this figure, only data for images that had at least five responsive cells were used.
**d**. Relationship between reverse filter correlation and noise correlation.
**e**. Relationship between signal and noise correlations.
**f**. Relationship between reverse filter correlation and signal correlation. (**d–f**) Data of responsive cell pairs were included. Colour indicates the number of cell pairs in each bin (bin size: 0.01).
**g–i.** Effect of noise correlation on image reconstruction performances (R in **g** and CD in **h**) and across-trial variability (**i**). Shuffle: trial-shuffled data. *: $p = 1.8\times10^{-5}$ (**g**), $p= 1.8\times10^{-5}$ (**h**) and $p=1.8\times10^{-5}$ (**i**) by signed rank test (N = 24 planes).
**j**. Distribution of noise correlation from all responsive cell pairs (0.03 [0.009–0.06]) in awake mice.
**k** and **l**. Distribution of noise correlation grouped by overlap index (**k**) or reverse filter correlation (**l**). Same as in **d–f** except for awake data.
**m–o**. Relationship among reverse filter correlation, noise correlation, and signal correlation in awake mice. Same as in **d–f** except for awake data and bin size (0.02).
**p–r.** Effect of noise correlation on image reconstruction performances (**p** and **q**) and across-trial variability (**r**) in awake mice. *: p = 0.03 (**p**), p= 0.047 (**q**) by signed rank test (N = 7 planes).

**Supplementary Figure 8. Response properties, response prediction and image reconstruction in awake mice (related to Fig. 1–5)**

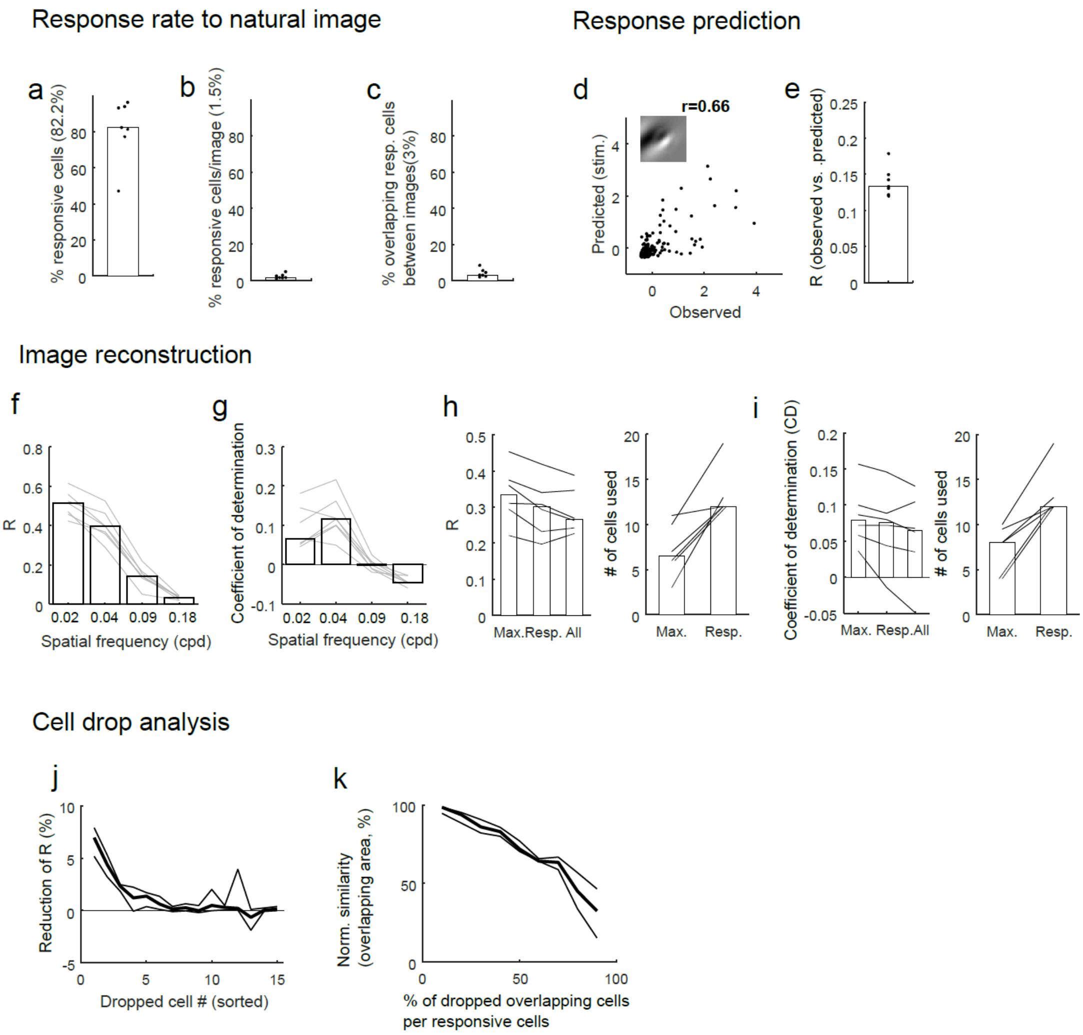


**Supplementary Figure 8. Response properties, response prediction and image reconstruction in awake mice (related to Fig. 1–5)**

**a**–**c**. Response properties (related to Fig. 1 in the main text).
The percentage of responsive cells (**a**), the percentage of responsive cells in each image (**b**), and the percentage of responsive cells that overlap between two images (**c**). Most neurons in a population were visually responsive to at least one image, whereas only a small number of neurons responded to each image. Each dot indicates one plane (n = 7 planes).
**d** and **e**. Response prediction (related to Fig.2). (**d**) An example of the response prediction of one cell. (**e**) Response prediction performance (n = 7 planes).
**f**–**i**. Image reconstruction performance by the cell-selection model (related to Fig. 3).
(**f** and **g**) Reconstruction performance in each spatial frequency (**f**: R and **g**: CD).
(**h** and **i**) Left panels: Performances obtained from all cells (All), responsive cells (Resp.) and the peak performance (Max.) were compared. The peak performance was detected within the number of responsive neurons. No significant difference was obtained in any pairs (signed rank test with Bonferroni correction). Right panels: The number of cells used for the reconstruction of each image in Resp. and Max. Peak performances were obtained from fewer cells compared to the number of responsive cells (**h**: P = 0.03; **i**: P = 0.03 by signed-rank test). In the analyses (**h** and **i**), we used only

data for images that had at least 10 responsive cells and excluded one plane of data because of a small number of samples.
**j** and **k**. Cell drop analysis (related to Fig. 5). (**j**) Reduction in R after a single-cell drop. The order of dropped cells was sorted using their evoked response amplitude (in descending order). A single-cell drop only slightly reduced the performance. (**k**) Reduction in reconstructed performance during the sequential drop of overlapping cells. The performance was estimated using the similarity (pixel-to-pixel correlation) of the overlapping area (see Fig. 5c, Supplementary fig. 6 and Methods). The analyses in (**j** and **k**) included only data for images which had at least five responsive cells.

**Supplementary Figure 9. Reliable image representation across trials in awake mice (related to Fig. 6)**

.

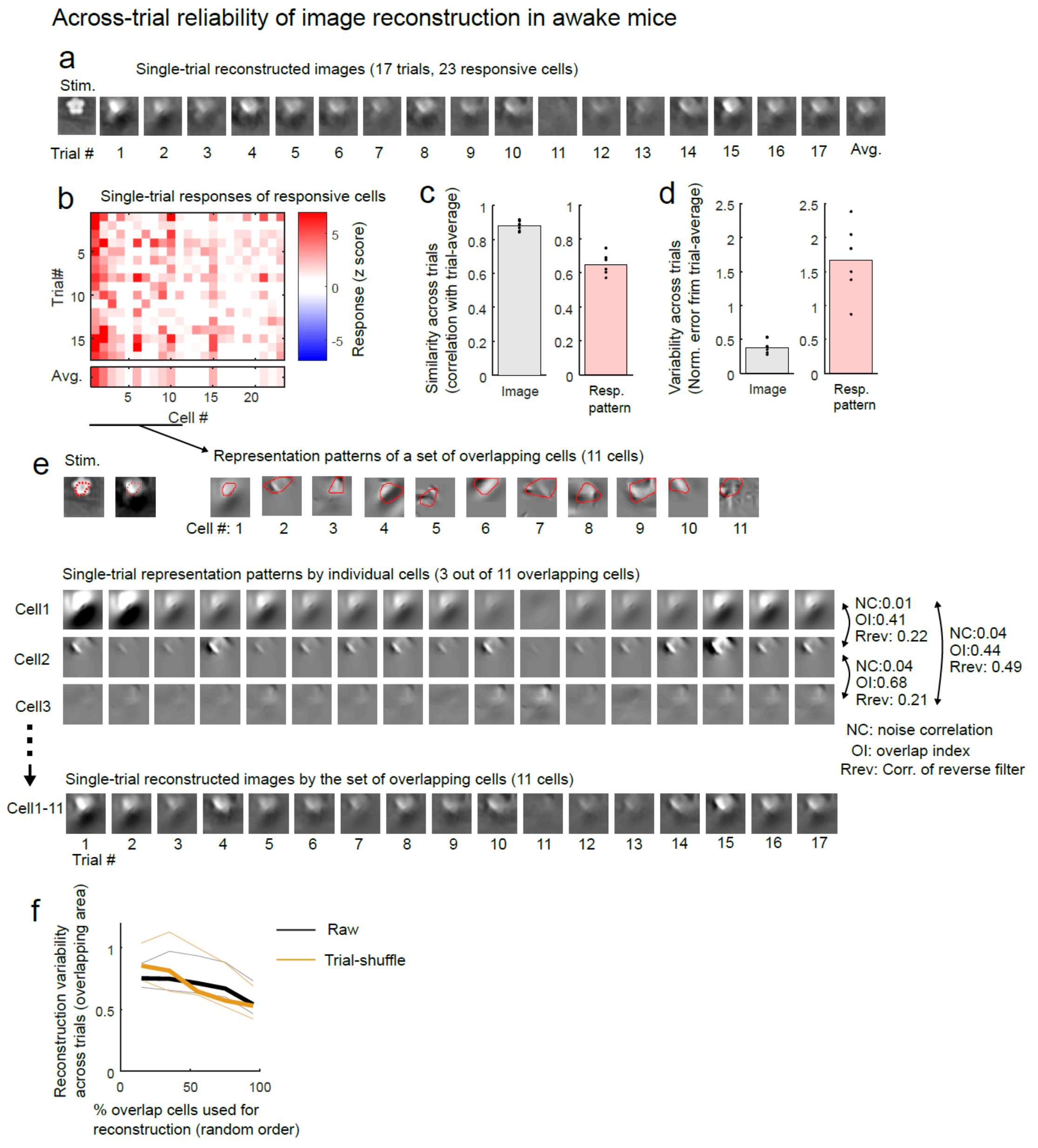


**Supplementary Figure 9. Reliable image representation across trials in awake mice (related to Fig. 6).**

**a**. Examples of single-trial reconstructed images. First panel: Stimulus image. Last panel: Trial-averaged image.
**b**. Single-trial evoked responses to the image in (**a**).
**c**. Across-trial similarity of reconstructed image (left) and of response pattern of the responsive cells (right). The across-trial similarity was a Pearson's correlation between a single-trial reconstructed image and trial-averaged image (N = 6 planes).
**d**. Across-trial variability of reconstructed images (left) and response patterns of responsive cells (right). Normalised squared error between single-trial image (or response patterns) and trial-averaged

image (or response patterns) was computed and used for the across-trial variability. (N = 6 planes).
**e**. Reconstructed image from a set of overlapping cells. Upper left panels: Stimulus and trial-averaged reconstructed images from the overlapping cells. The red dotted line indicates the overlapping area. Upper right panels: Representation (reverse filters) of the overlapping cells. The red line in each panel indicates the representation area. Cell 1 was the reference cell. Lower panels: Single-trial representation of three example cells selected from the overlapping cells. Bottom panels: Single-trial reconstructed image obtained from the overlapping cells.
**f**. Across-trial variability against the percentage of the overlapping cells used for the reconstruction. N = 6 planes. Black lines: Raw data. Orange lines: trial-shuffled data. Thick and thin lines were median and $25^{th}$ or $75^{th}$ percentile of the variability among 6 planes.
In the analyses shown in this figure, we used only data for images that had at least five responsive cells and excluded one plane of data because of the small number of samples.

**Supplementary Figure 10. Image representation in excitatory and inhibitory cells.**

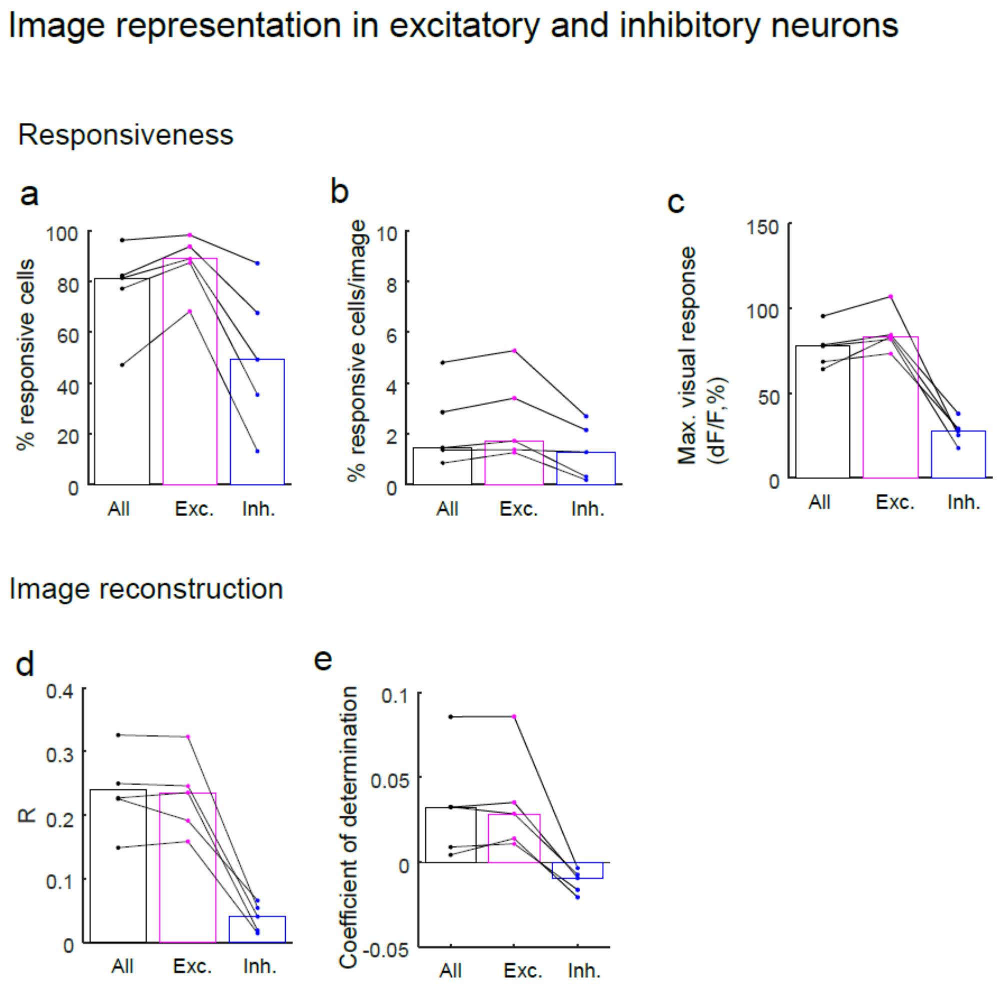


**Supplementary Figure 10. Image representation in excitatory and inhibitory cells.**

**a**–**c**. Response properties in excitatory (Exc.), inhibitory (Inh.), and all cells including both types (All). (**a**) The percentage of responsive cells. (**b**) The percentage of responsive cells for each image. (**c**) Maximal evoked response.

**d** and **e**. Image reconstruction performances. The performances of excitatory cells were almost comparable with those of all cells, indicating that excitatory cells mainly represented the images. (**a**–**e**) N = 5 planes.

**Supplementary Figure 11. Comparison of image reconstructions between staying and running periods**

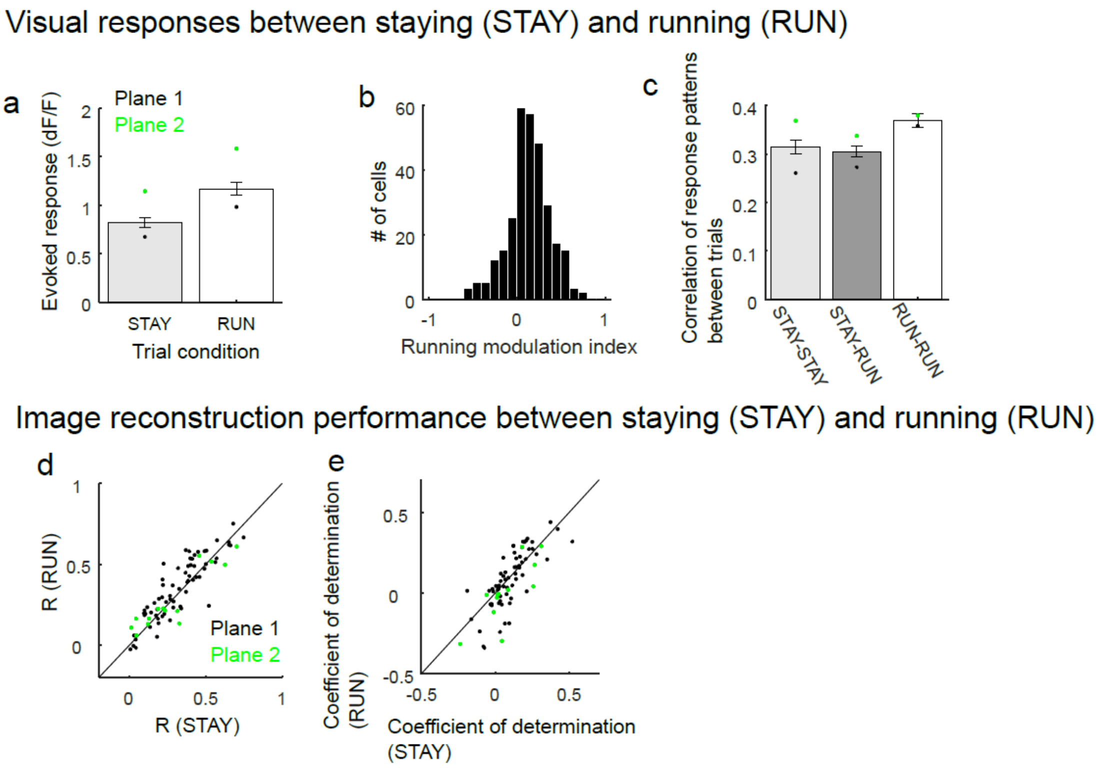


**Supplementary Figure 11. Comparison of image reconstructions between staying and running periods**

This analysis used only data for images that had at least 5 responsive cells and 4 trials of both staying and running conditions (data for 80 image cases, 295 responsive cells from two planes).
**a**–**c**. Comparisons of visual responses between staying and running periods. (**a**) Evoked response amplitude (n = 295 responsive cells). Bars and error bars indicate the mean and standard error of means across all cells used in this analysis. (**b**) Distribution of the running modulation index (n = 295 cells). (**c**) Correlation coefficients of response patterns between trials (n = 80 image cases). Although visually evoked responses during running tended to be higher than those during staying, response patterns were similar between the conditions. (**a** and **c**) Each dot indicates the mean of each plane.
**d** and **e**. Comparison of image reconstruction performance between the conditions. (**d**) Pixel-to-pixel correlation between stimulus and reconstructed images (R). Each dot indicates each image case (STAY: 0.31±0.005. RUN: 0.34±0.005, p = 0.02 by signed-rank test, n = 80 images). (**e**) Coefficient of determination (CD. STAY: 0.08 ± 0.004. RUN: 0.03 ± 0.007, p = 0.07 by signed-rank test, n = 80 images). Different colours of dots indicate data from different planes.
